# Illuminating microbial phosphorus cycling in deep-sea cold seep sediments using protein language models

**DOI:** 10.1101/2024.07.09.602434

**Authors:** Chuwen Zhang, Yong He, Jieni Wang, Tengkai Chen, Federico Baltar, Minjie Hu, Jing Liao, Xi Xiao, Zhao-Rong Li, Xiyang Dong

## Abstract

Phosphorus is essential for life and critically influences marine productivity. Despite geochemical evidence of active phosphorus cycling in deep-sea cold seep ecosystems, the microbial processes involved remain poorly understood. Traditional sequence-based searches often fail to detect proteins with remote homology. To address this, we developed a protein language model (PLM) named LucaPCycle that integrates sequence and structural information. This PLM-based approach identified 4,606 new phosphorus-cycling protein families based on the non-redundant gene and genome catalogs from global cold seeps, substantially enhancing our understanding of their diversity, ecology, and function. Among previously unannotated sequences, we discovered three novel alkaline phosphatase families (ALP1, ALP2 and ALP3) that feature unique domain organizations and preserved enzymatic capabilities. These results highlight previously overlooked ecological importance of phosphorus cycling within cold seeps, corroborated by data from porewater geochemistry, metatranscriptomics, and metabolomics. We identified a previously unrecognized diversity of archaea contributing to organic phosphorus mineralization and inorganic phosphorus solubilization through various mechanisms. This includes ecologically significant groups such as Asgardarchaeota, anaerobic methanotrophic archaea (ANME), and Thermoproteota. Additionally, viruses can enhance their hosts’ (e.g., ANME) phosphorus utilization through the PhoR-PhoB regulatory system and PhnCDE transporter, indirectly influencing methane dynamics. Overall, our PLM-based functional predictions are capable of accessing previously ’hidden’ sequence spaces for microbial phosphorus cycling, and can be applied to other various ecosystems.

## Introduction

Cold seeps, often situated along continental margins, are areas where hydrocarbon-rich fluids escape from subsurface reservoirs to the seafloor^1, 2^. These unique environments are distinguished by their chemosynthetic microorganisms that utilize methane and other hydrocarbons—such as non-methane alkanes and aromatic hydrocarbons—as sources of carbon and energy to sustain life^3–5^. A key process in these habitats is the anaerobic oxidation of methane (AOM), which occurs in conjunction with sulfate reduction, mediated by a consortium of anaerobic methanotrophic archaea (ANME) and sulfate-reducing bacteria (SRB)^6, 7^. In addition to carbon and sulfur, these microbial communities also require phosphorus for essential biological functions such as building cell membranes, synthesizing nucleic acids, and producing energy carriers and various phosphorylated metabolic intermediates^8, 9^. Despite its importance, studies on phosphorus cycling in deep-sea cold seep sediments are less developed than those on carbon and sulfur^3, 10^.

Chen et al. (2023) examined a piston core from the Haima cold seeps in the Qiongdongnan Basin, northern South China Sea, revealing high phosphate concentrations in porewater and low ratios of organic phosphorus to total organic carbon (TOC)^11^. This suggests a phosphorus-rich environment within these deep-sea cold seeps. The study, along with others, provided geochemical evidence that AOM is a key driver of phosphorus cycling within these environments^11–13^. This process, associated with the generation of hydrogen sulfide, leads to the reductive dissolution of iron oxides, releasing iron-bound phosphorus into the surrounding porewater. Another study indicates that in deep-sea carbonate deposits from cold seeps in the South and East China Seas, calcium-bound phosphorus predominates, co-precipitating with calcium carbonate during AOM, with iron also playing a crucial role in the release of phosphorus from iron oxides^14^. These findings underscore the complex interactions between microbial activity and chemical reactions in deep-sea phosphorus cycling, through a combination of microbial and chemical processes. However, the specific contributions of microbially-driven processes to the production of dissolved inorganic phosphorus in these environments remain unclear.

Microbes have evolved various enzymatic mechanisms to convert recalcitrant phosphorus forms into bioavailable phosphorus^15^. These phosphorus-cycling microorganisms (PCMs) possess genes related to four main processes: organic phosphorus mineralization, inorganic phosphorus solubilization, phosphorus uptake and transport, and phosphorus-starvation response regulation^16–19^. Organic phosphorus mineralization primarily involves release of hydrolytic enzymes to recycle various organic phosphorus compounds. Acid (e.g., PhoN, PhoC) and alkaline phosphatases (e.g., PhoD, PhoA) cleave phosphate from complex phosphoesters^20–22^, while phytases (e.g., AppA) hydrolyze phosphate from phytate or myo-inositol hexakis dihydrogen phosphate^23^. Phosphonatase (e.g., PhnW, PhnX) and C-P lyase (PhnGHIJKLM) participate in C-P cleavage of phosphonates^24, 25^. For inorganic phosphorus mobilization, polyphosphate kinase (PPK) catalyzes the formation of polyphosphate (poly-P), and exopolyphosphatase (PPX) subsequently releases phosphate from poly-P chains^26, 27^. PCMs also secrete organic acids, such as gluconic acid produced by the oxidation of glucose via membrane-bound quinoprotein glucose dehydrogenase (Gcd), to solubilize inorganic phosphorus minerals^28^. Additionally, PCMs harbor high-affinity (PstABCS) and low-affinity (Pit) transporters for phosphate uptake under varying conditions^29^, and can adjust phosphorus-sensing and regulatory mechanisms (e.g., PhoB-PhoR system) to optimize phosphorus use under scarcity^19, 30^.

The annotation of phosphorus-cycling genes has traditionally relied on sequence homology methods such as Blastp or profile hidden Markov models (pHMMs)^30, 31^. However, these methods are limited to detecting sequences with relatively high similarity, potentially missing divergent sequences with distant evolutionary relationships. To overcome this limitation, researchers are increasingly using protein structure features, which are more conserved over evolutionary timescales and crucial for maintaining functional integrity^32^. Recent advancements in deep learning for protein bioinformatics, exemplified by tools like AlphaFold, have facilitated insights into these previously obscure ’functionally dark’ proteins^33–35^. For example, this approach led to the discovery of a new superfamily of translation-targeting toxin-antitoxin systems, TumE–TumA^34^. Additionally, the field of functional annotation predictions has benefited significantly from the application of natural language processing (NLP) techniques. In NLP, protein language models (PLMs) are developed using extensive labeled and unlabeled datasets to create vectorized embeddings that represent both protein sequences and structures. This innovative method has been applied in various life science research areas, including the identification of new viruses, the discovery of antibiotics, and the detection of remote proteins^32, 36–38^.

In this study, we aim to improve the annotation of phosphorus-cycling proteins, focusing on previously unexplored sequence spaces, to deepen our understanding of microbial contribution in phosphorus cycling across global cold seeps. To achieve this, we developed a novel protein language model named LucaPCycle, specifically designed to target phosphorus-cycling proteins. This model was trained using both protein sequence and structural data. Our findings indicate that LucaPCycle effectively identifies remote homologies, thus providing a valuable complement to traditional alignment-based methods. This enhancement allows for a more comprehensive understanding of the roles and mechanisms of phosphorus cycling, not limited to the cold seep ecosystems investigated here, but also in other environments.

## Results and discussion

### PLM-based LucaPCycle outperforms homology-based approaches

LucaPCycle is a dual-channel deep learning model (**Fig. 1a**). The first channel utilizes the protein language model ESM2 to extract features from sequences at the residue level, leveraging its self-supervised learning capabilities to understand the sequence context^39^. The second channel involves a Transformer-Encoder^40^ designed to capture the features of the raw sequence. We applied LucaPCycle to build two models, one for binary-classification and the other for 31-classification. The binary-classification model predicts whether a protein sequence has a phosphorus-cycling function. If it does, the 31-classification model then categorizes it into one of 31 specific types of phosphorus-cycling genes (detailed in **Table S1**). LucaPCycle is established based on 214,193 positive samples of 31 phosphorus-cycling proteins and 751,331 negative samples covering a range of functions, including intracellular phosphorus metabolism, nitrogen cycling, sulfur cycling, and other unrelated functions **(Fig. S1a-S1b**). Detailed information on model building is provided in the **Methods** section.

**Figure 1.**
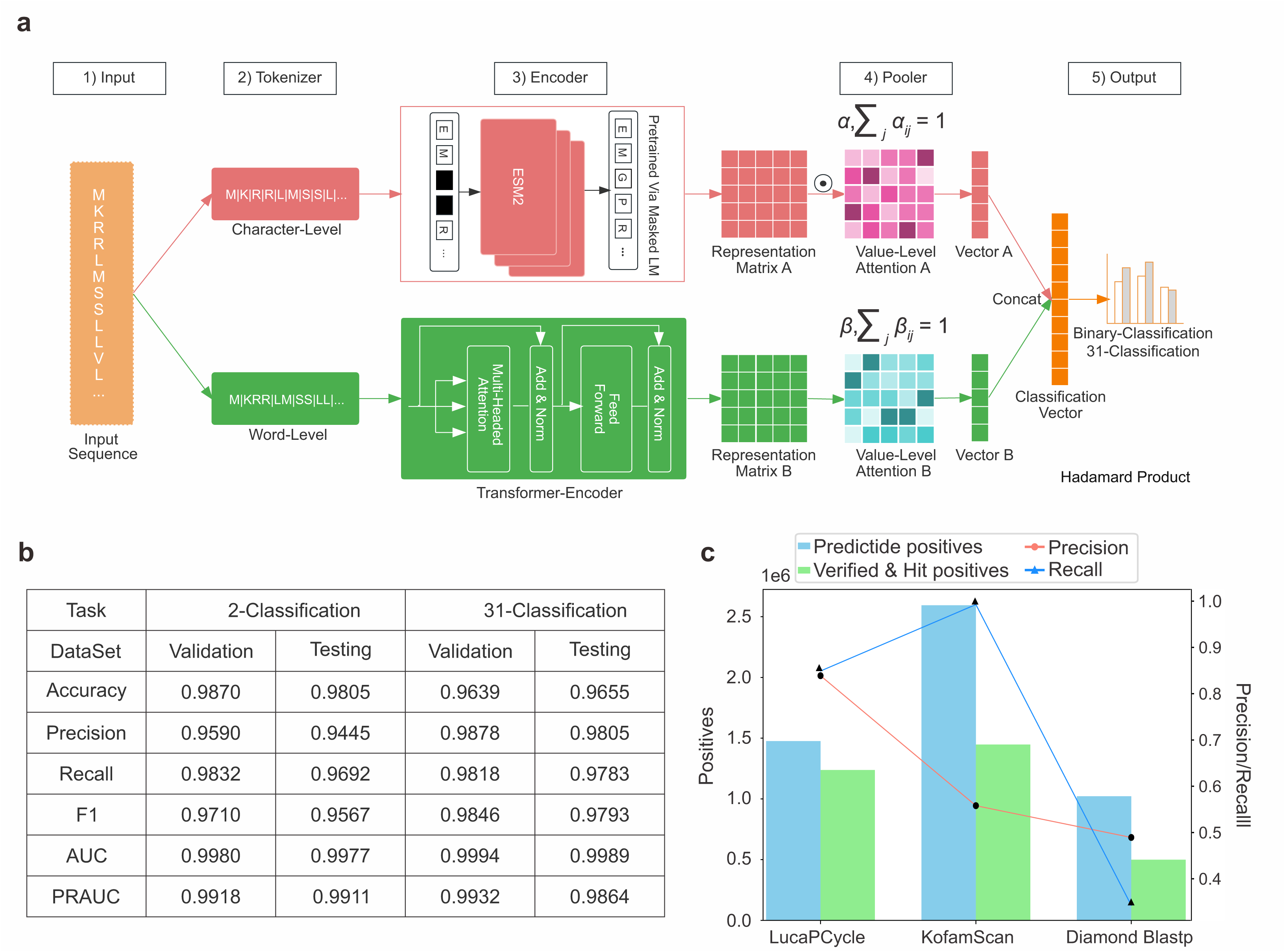
Overview of LucaPCycle for phosphorus-cycling protein annotation. **a**, Architecture of LucaPCycle. LucaPCycle consists of five modules: Input, Tokenizer, Encoder, Pooler, and Output. The Tokenizer module performs character-level and word-level tokenization on input sequences. The Encoder automatically extracts two types of features: the representation matrix of the protein language model and the representation matrix of the raw sequence. The Pooler selects essential features and transforms the feature matrix into a vector. The Output layer concatenates these two pooled vectors for classification. **b**, Performance of binary-classification and 31-classification models within LucaPCycle. Six metrics including accuracy, precision, recall, F1-score, AUC, and PRAUC were evaluated based on the validation and test datasets. **c**, Benchmarking of LucaPCycle with KofamScan and Diamond Blastp.

Binary- and 31-classification models in LucaPCycle both perform well on metrics of accuracy, precision, recall, F1-score (The harmonic mean of the precision and recall), AUC (Area under the ROC curve), and PRAUC (Area under the precision-recall curve), achieving scores above 0.94 when evaluated on validation and test datasets (**Fig. 1b**). To further demonstrate the sensitivity and specificity of LucaPCycle, we benchmarked it to two homology-based functional annotation tools, utilizing a dataset of 1,463,634 true positive sequences (see **Methods**). The homologous protein search methods include: (1) Diamond Blastp based on sequence similarity with curated phosphorus-cycling genes in PCycDB database^30^; and (2) KofamScan^31^, a gene function annotation tool based on KEGG Orthology and hidden Markov model (HMM). Notably, LucaPCycle outperforms the other two methods with the highest precision rate (84%, a measure of the accuracy of the positive predictions made by the model) and a comparable recall rate (85%, a measure of how well the model can identify all the positive samples; **Fig. 1c**). In comparisons, KofamScan exhibited a high recall rate (99%) but a low precision rate (56%), while Diamond Blastp exhibited both the lowest precision rate (49%) and recall rate (34%).

### Remote homology detection: three novel alkaline phosphatases at cold seeps

Using the non-redundant gene (n = 147,289,169) and genome (n = 1,878) catalogs from global cold seeps, phosphorus-cycling genes were functionally annotated using LucaPCycle and two sequence homology-based methods, Diamond Blastp and KofamScan (**Fig. S2**). The results from these three search strategies were cross-checked by protein domain analysis, a deep learning model DeepFRI^41^ and a machine learning model CLEAN^42^. Subsequently, all validated phosphorus-cycling proteins were merged and clustered using sequence-based clustering algorithm to generate representative sequences for phylogenetic analyses. Singletons, likely indicative of sequencing errors^43^, were discarded, leading to the identification of 321,789 non-singleton phosphorus-cycling protein families (**Fig. S2**). These families predominantly consisted of sequences organized in small groups (2 members) and exhibited diverse lengths with the most common range being 100 to 300 amino acids (**Fig. S3**). Among 321,789 non-singleton phosphorus-cycling protein families, 4,606 phosphorus-cycling protein families were exclusively identified by LucaPCycle, possibly representing proteins with distant evolutionary relationships (**Fig. S4a**). These remote sequences cover 26 distinct types of phosphorus-cycling genes, with the highest number of sequences being *phnC* (n = 18,280), *pstB* (n = 17,846), *phoB* (n = 13,782), and *ppx* (n = 12,050), respectively (**Fig. S4b**). These remote proteins could significantly contribute to the biogeochemical cycling of phosphorus in cold seeps, but have been previously neglected due to limitations of homology-based methods. Notably, the *ppx* gene with remote homology constituted substantial proportions of the total *ppx* gene (75%) and transcript (74%) abundances (**Fig. S4c-S4d**).

We further explored the remote homology information of alkaline phosphatases (ALPs) recovered by LucaPCycle. This group was chosen due to its prevalence in global ocean water and its clinical significance^20, 44, 45^. ALPs are typically separated into three classical families (PhoA, PhoD, and PhoX)^20, 22^ and one divergent family (PafA)^46^. Our sequence and structure tree topologies showed that remote ALPs formed three distinct clusters (**Fig. 2a-2b**). We designated these clusters as ALP1, ALP2, and ALP3 in this study. Specifically, sequence and structural phylogenies placed ALP1 between PhoX and divergent PafA, while ALP2 and ALP3 were positioned closer to PafA and PhoA, respectively. Despite sharing a low sequence identity averaging at 14.3%, a large proportion of ALP1 demonstrated structural homology to classic PhoD (TM-score > 0.5; **Fig. S5a**). ALP2 were more resemblance to divergent PafA rather than to other three classic ALPs. All ALP2 displayed a high structural similarity with divergent PafA, with the majority exceeding a TM-score of 0.75 but sequence similarity averaged at 33.6% (**Fig. S5b**). ALP3 shared a sequence identity of 17.6% with PhoA from *Escherichia coli* (PDB: 1AJA) but exhibited a TM-score of 0.82 in terms of structure (**Fig. S5c**). Thus, ALP1, ALP2, and ALP3 could be remote relatives of PhoD, PafA and PhoA, respectively.

**Figure 2.**
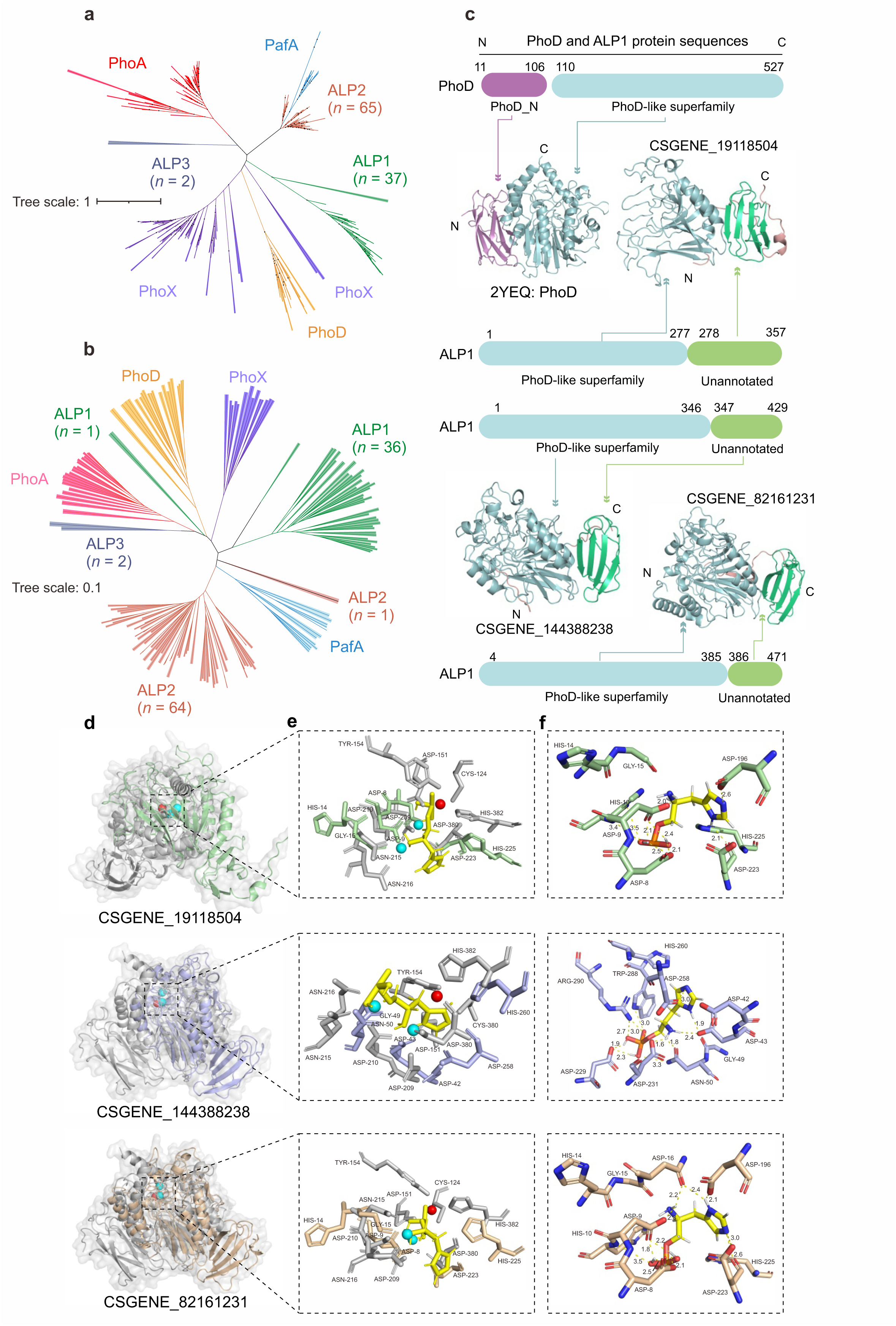
Three novel alkaline phosphatases (ALPs) with remote homology identified by LucaPCycle. **a**, Phylogenetic tree of ALPs based on protein sequences. Scale bar indicates the mean number of substitutions per site. Bootstrap values over 70% were shown. The n values denote the count of ALPs sequences within each family. **b**, Structure-based phylogeny of ALPs. All protein 3D structures are predicted by AlphaFold3. **c**, Comparison of domains between ALP1 and classic PhoD identified by sequence (InterproScan) versus structure-based methods (ECOD). **d**, Structural comparison between three predicted remote ALP1 and the experimentally verified PhoD (2YEQ, in grey). **e**, Close-up view of the putative active sites for metal binging. The active site residues of PhoD are shown in grey sticks. The catalytic site metal ions are presented as spheres, with Fe^3+^ shown in red and Ca^2+^ shown in cyan. **f**, Molecular docking of ALP1 and phosphomonoesters. Substrates histidinol phosphate is shown in yellow sticks.

Protein structures are composed of domains, independently folding units that can be found in multiple structural contexts and functional roles^47^. Extracting these domains enables a more fine-grained description of the features among different proteins. Subsequently, we compared the domain variations between ALP1 and the classic PhoD, given their relatively low TM-score (**Fig. S5a**). Both ALP1 and PhoD were partitioned into two domains by sequence- and structure-based domain assignments (**Fig. 2c**). However, their domain organizations were completely different. Typically, PhoD features a core architecture consisting of a sandwich of long β-sheets flanked by α-helices of varying lengths^48^ and an upstream N-terminal β-sheet domain. In contrast, ALP1 lacked the N-terminal domain and was organized with a core architecture followed by an unannotated C-terminal β-sheet domain (**Fig. 2c**). ALP1 also exhibited fewer α-helices in its core architecture compared to PhoD. However, the core architectures of ALP1 and PhoD maintained a similar orientation and were conserved within the ECOD hierarchy at various levels (**Fig. 2d and Table S2**): A-group (same architecture), X-group (possible homology), H-group (homology), T-group (topology), and F-group (sequence similarity). Nevertheless, the C-terminal domain of ALP1 and the N-terminal domain of PhoD were structural divergence and assigned different ECOD T-group and F-group labels (**Fig. 2d and Table S2**). The emergence of a C-terminal β-sheet domain may represent an adaptive strategy for phosphorus-cycling microbes to enhance fitness in extreme deep-sea environments.

Despite evolutionary variations in amino acid sequences and protein structures, the remote ALP1 retained the function of phosphoester hydrolysis. ALP1 preserved metal binding sites in their core architectures, with active center residues exhibiting similar spatial distribution to classic PhoD (**Fig. 2e**). The molecular docking analysis demonstrated robust and stable interactions between ALP1 and 89 various phosphoester compounds, as indicated by low binding energies (-11.02∼-2.39 kcal/mol; **Fig. 2f and Table S3**). This conservation of essential domains (i.e., core architectures) for function maintenance in remote ALPs suggests convergent evolution.

### Phosphorus-cycling genes are diverse, active and depth-stratified at cold seeps

The microbes inhabiting cold seeps were found to harbor multiple phosphorus acquisition strategies. In particular, genes encoding the regulatory system (PhoR-PhoB) were among the most predominant (av. 3741.6 genes per million, GPM; **Fig. 3a**). These microbes also exhibited efficient phosphate transport systems to acquire available phosphorus with minimal energy investment. This was evidenced by the significantly higher levels of genes encoding for high-affinity phosphate transporter (PstABCS; av. 533.8 GPM) compared to those encoding the low-affinity phosphate transporter (Pit; av. 148.8 GPM). The prevalence of PhoR-PhoB and PstABCS aligned with observations in soil microbes in response to limited bioavailable phosphorus^49, 50^. Within inorganic phosphorus solubilization, genes encoding glucose dehydrogenase (Gcd), polyphosphate kinase (PPK) and exopolyphosphatase (PPX) demonstrated comparable abundance patterns.

**Figure 3.**
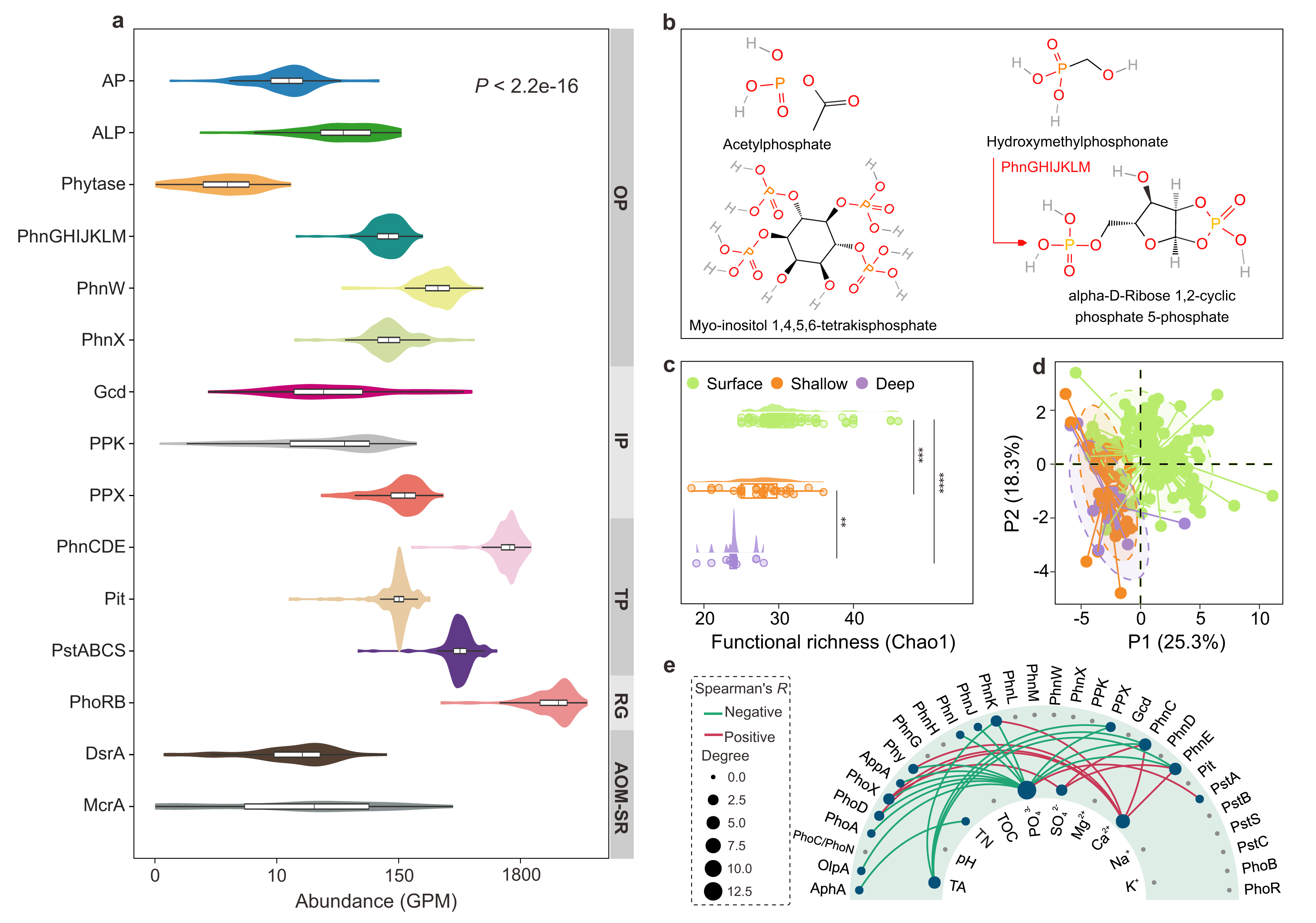
The biogeographic pattern of phosphorus-cycling genes across global cold seeps. **a**, Relative abundances of phosphorus-cycling genes from four distinct metabolic processes, as compared to *mcrA* and *dsrA*. OP: Organic phosphorus mineralization; IP: Inorganic phosphorus solubilization; TP: Transporters; RG: Regulatory genes; AOM-SR: The anaerobic oxidation of methane (AOM) coupled to sulfate reduction (SR). ALP: alkaline phosphatase including PhoA, PhoD and PhoX. Acid phosphatase including AphA, OlpA, PhoN and PhoC. Gene abundances are indicated as GPM (genes per million). *P* values of differences among different genes were calculated using Kruskal-Wallis rank-sum tests. **b**, The chemical structures of phosphomonoester, phytate, phosphonates and metabolic intermediates identified in sediments from Qiongdongnan and Shenhu cold seeps. **c**, Comparison of functional richness of phosphorus-cycling genes across diverse cold seep sediment depths. Surface, < 1 mbsf; Shallow, 1 to 10 mbsf; Deep, > 10 mbsf. Asterisks indicate statistically significant differences among groups of surface, shallow, and deep sediments (determined by two-sided Wilcoxon rank sum test; \*\**P* < 0.01 and \*\*\**P* < 0.001). **d**, The partial least squares discrimination analysis (PLS-DA) plots depicts the differences in phosphorus-cycling microbial communities among different sediment depths. Analysis of similarity were examined using a 999-permutation PERMANOVA test. **e**, Spearman correlation between the abundances of phosphorus-cycling genes and various physicochemical variables. The red and blue colors indicate positive and negative correlations, respectively. TA: total alkalinity; TOC: total organic carbon; TN: total nitrogen.

Organic phosphorus constitutes about 25% of the total phosphorus buried in marine sediments^51^, where phosphorus atoms are covalently bonded to carbon either directly (P–C, phosphonates) or through a phosphoester linkage (P–O–C, phosphomonoesters, phosphodiesters, phosphotriesters and phytates)^52^. Depending on the substrate, microorganisms rely on distinct enzymes to mineralize organic phosphorus and release inorganic phosphate. For phosphoester hydrolysis, genes for nonspecific ALPs (*phoA*, *phoD*, or *phoX*, av. 58.1 GPM) were more prevalent than genes for nonspecific acid phosphatases (APs; *aphA*, *olpA*, *phoN* or *phoC*, av. 16.1 GPM) and phytases (av. 4.0 GPM; **Fig. 3a**). The lower abundance of genes encoding phytase can be attributed to the limited availability of its substrate, phytates, sourced primarily from terrestrial runoff ^53^. Additionally, the approximately neutral pH at cold seeps may limit the activity of APs, which has an optimal pH range of 4–6^54^ (**Table S4**). Regarding phosphonate degradation, the genes encoding substrate-specific phosphonatase (PhnW-PhnX) and broad-specificity C–P lyase (PhnGHIJK) showed comparable abundances (**Fig. 3a**). Moreover, untargeted metabolomics detected over 104 organic phosphorus compounds in 55 cold seep sediments (**Fig. 3b and Table S5**). These metabolites included a wide diversity of phosphomonoesters (e.g., acetylphosphate), phytates (e.g., 1D-Myo-inositol 1,4,5,6-tetrakisphosphate) and phosphonates (e.g., Hydroxymethylphosphonate). Additionally, our results identified intermediate metabolites of organic phosphorus mineralization, such as alpha-D-ribose 1,2-cyclic phosphate 5-phosphate, generated during phosphonate degradation. This provides potential biological evidence for the recycling of organic phosphorus molecules at cold seeps. However, it cannot conclusively rule out the possibility that some of these molecules originate from external sources.

To determine the biogeographic patterns of phosphorus-cycling proteins, each metagenome was categorized by its sediment depth (i.e., surface: < 1 mbsf; shallow: 1–10 mbsf; deep: > 10 mbsf). The distribution of phosphorus-cycling proteins was stratified by sediment depth. The functional richness (Chao1) decreased with depth and peaked at surface layers (*P* < 0.001; **Fig. 3c**). The compositions of phosphorus-cycling proteins also displayed significant variations among surface, shallow, and deep layers (*F* = 13.2, *P* = 0.001, *R*^2^ = 0.14, 999-permutations test; **Fig. 3d**). Likewise, we observed depth-related pattern in the expression of phosphorus-cycling genes, with a higher transcript abundance detected in surface layers (averaging from 1.3 to 1244.9 TPM; **Fig. S6**). Measurements of pore water samples from 12 sediment cores in the Shenhu area and the Qiongdongnan basin revealed that PO ^3−^ concentration increases with sediment depth (**Fig. S7 and Table S6**). Additionally, PO_4_ ^3−^ concentration exerted a negative feedback regulation on the abundance of phosphorus-cycling genes (**Fig. 3e**), as observed in soil bacteria^55^. Thus, this depth effect can be driven by PO_4_^3−^ concentrations. Other factors, including cold seep types (i.e., gas hydrate, methane seep, oil and gas seep, mud/asphalt volcano), also contribute to the patterns of phosphorus-cycling gene abundance and expression (**Figs. S6 and S8**). Significant differences in functional richness (*P* < 0.001; **Fig. S8a**) and compositions of phosphorus-cycling proteins among cold seep types were observed (*F* = 6.78, *P* = 0.002, *R*^2^ = 0.11, 999-permutations test; **Fig. S8b**).

Subsequently, we assessed the importance of phosphorus cycling in cold seep ecosystems based on the abundance of corresponding genes and transcripts. We chose *mcrA* and *dsrA* genes as references due to their key roles in anaerobic methane oxidation coupled with sulfate reduction, a major biological process in cold seeps^56, 57^. Except for genes encoding APs and phytases, the abundances of other phosphorus-cycling genes associated with 11 distinct metabolic processes (averaging from 58.1 to 3741.6 GPM) exceeded those of *mcrA* (av. 46.1 GPM) or *dsrA* genes (av. 21.3 GPM; **Fig. 3a**). Likewise, the expression levels of phosphorus-cycling genes (averaging from 6.6 to 422.8 TPM) demonstrated a similar pattern when compared to *mcrA* (av. 251.5 TPM) or *dsrA* genes (av. 5.7 TPM; **Fig. S9**). To date, microbial transformation of phosphorus is well-documented in ocean surface water where phosphate is often limited^24, 44^. Here, our results demonstrated that phosphorus cycling driven by microorganisms is clearly important within deep-sea cold seeps in addition to the extensively studied carbon and sulfur metabolisms^56, 57^.

### Archaea are key players for organic phosphorus mineralization and inorganic phosphorus solubilization

Among non-singleton phosphorus-cycling protein families, 1,820,568 sequences were identified in Bacteria and 149,641 in Archaea, leaving 162,460 as unclassified (**Fig. S10**). The subsequent analysis was focused on the cold seep genome catalog to ensure a finer taxonomic assignment of phosphorus-cycling proteins. The results revealed that genes encoding the regulatory system PhoR-PhoB, the high-affinity PstABCS and the low-affinity Pit were widely distributed throughout all major archaeal and bacterial phyla (**Fig. S11**). This broad distribution pattern highlights the foundational role of phosphorus uptake and regulation in supporting cell growth^9^. Aside from the aforementioned basic genetic equipment, our findings substantially expanded the taxonomic distribution of genes related to the phosphate scavenging from inorganic and organic phosphorus source among Archaea domain (**Fig. 4a**). To date, genes encoding inorganic polyphosphate (poly-P) metabolism, specifically PPK or PPX, have been identified in the archaeal phyla Halobacteriota and Thermoproteota^58^. However, our PLM-based LucaPCycle analysis alternatively identified PPX in Iainarchaeota (formerly Diapherotrites) from the DPANN superphylum. DPANN are characterized by small genome sizes and minimal metabolic function^59^. The selective maintenance of *ppx* genes in DPANN archaea may be associated with changing environments, as poly-P confers resistance to environmental stresses^60^. However, mineral phosphorus solubilization in archaea has received less attention with the majority of studies focusing on bacteria^17, 18, 28^. Beyond Halobacteriota^28^, four additional archaeal phyla—Hydrothermarchaeota, Aenigmatarchaeota, Thermoplasmatota, and Thermoproteota—also harbored the *gcd* gene, which encodes glucose dehydrogenase (**Fig. 4a**). This enzyme catalyzes the oxidation of glucose to gluconic acid, facilitating the solubilization of mineral-sorbed inorganic phosphate^18^.

**Figure 4.**
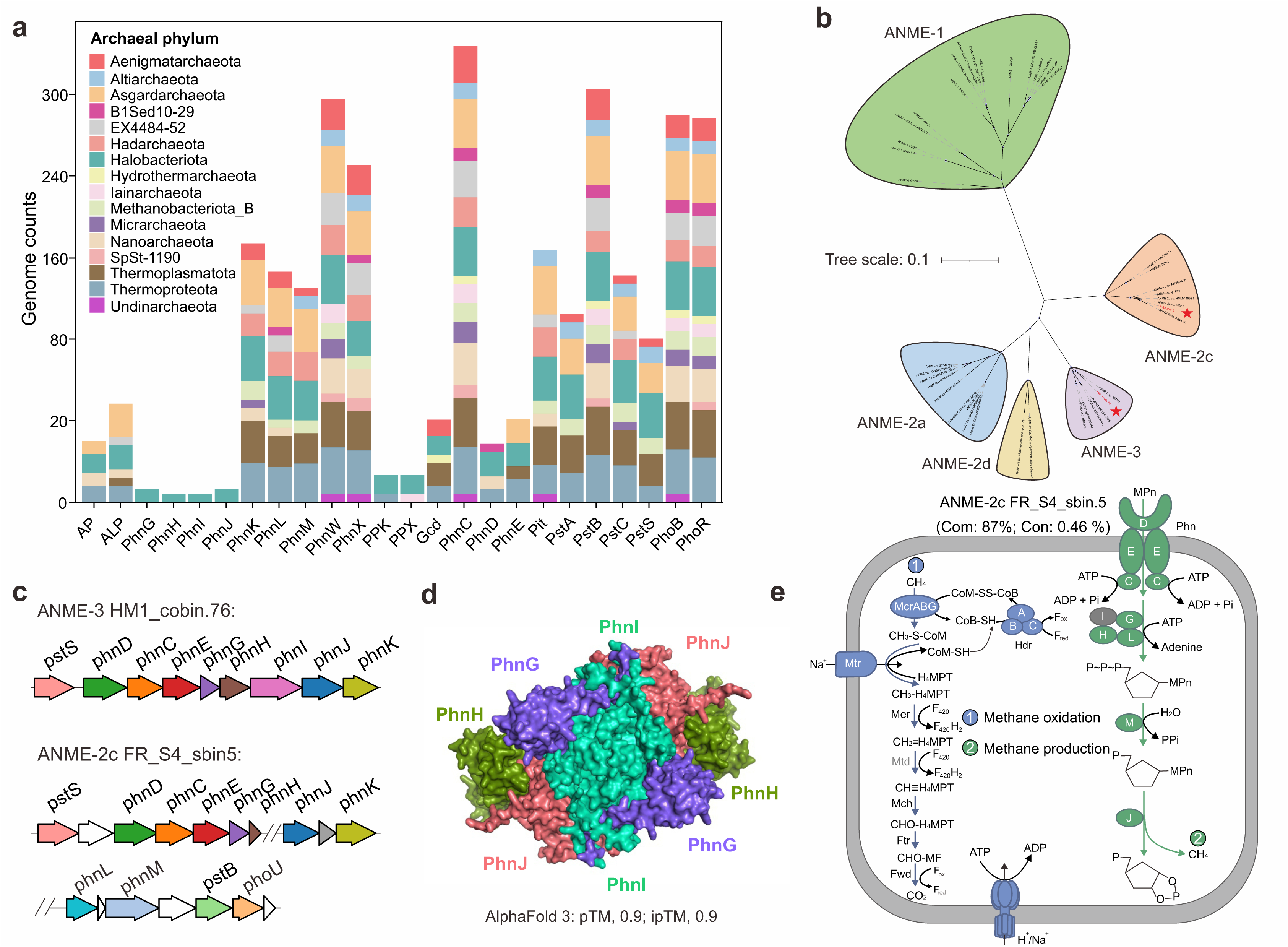
The overlooked role of archaea in deep-sea phosphorus-cycling. **a**, Phylogenetic distribution of phosphorus-cycling genes within Archaea domain. **b**, Taxonomic placement of PhnJ-containing archaeal genomes. Phylogenomic tree constructed based on alignments of 43 conserved protein sequences from MAG FR_S4_sbin5 and HM1_cobin.76 (indicated by red stars). Bootstrap value ≥ 70 are shown. Scale bar indicates amino acid substitutions per site. **c**, Organization of C-P lyase operons in ANME-2c and ANME-3. **d**, The structure of Phn(GHIJK)_2_ complex shown as a surface representation. The C-P lyase core complex and the interaction among subunits were predicted using AlphaFold3. **e**, Anaerobic oxidation of methane and methanogenesis in ANME-2c. Grey color indicates the absence of the enzyme. The percentages between brackets indicate the estimated completeness of the corresponding MAGs. MPn: methylphosphonate. A complete list of metabolic information can be found in **Table S14**.

Contrary to a recent study reporting that most archaea cannot metabolize phosphonates^61^, this study found that a variety of archaeal phyla possess the genetic capability to do so. Specifically, genes responsible for phosphonate substrate-specific catabolism (*phnWX*), which target 2-AEP, demonstrated a wide taxonomic distribution across archaea domain (**Fig. 4a**). These genes co-occurred in 12 different archaeal phyla, predominantly Halobacteriota, Thermoproteota, Asgardarchaeota, and Thermoplasmatota, whereas in global seawater, they were mainly found in the bacterial phylum Proteobacteria^24^. Additionally, the broad-specificity catabolism genes (C-P lyase, PhnGHIJKLM) that hydrolyze a broad range of phosphonates were identified in genomes affiliated with archaea Halobacteriota. Notably, metagenome-assembled genomes (MAGs; FR_S4_sbin5 and HM1_cobin.76) belonging to ecologically important methanotrophs ANME-2c and ANME-3 possessed near complete C-P lyase operon (**Fig. 4b-4c**). This marks a significant discovery, overturning the previous notion that the *phnJ* gene was exclusive to bacteria in marine ecosystems^24, 62, 63^. Organization of C-P lyase operons exhibited a consecutive string of C-P lyase subunits and alongside the synteny with other phosphorus-cycling genes, such as the phosphonate transporter (PhnCDE) and partial phosphate transporter (PstABCS). The neighborhood occurrence of these genes in C-P lyase operons reflects an evolutionary adaptation to enhance the efficiency of phosphorus-related mechanisms, consistent with what has been observed in bacteria^24^. The subunits of the C-P lyase core complex found in ANME archaea interacted closely (pTM: 0.9; ipTM: 0.9) to form conformational states necessary for phosphonate breakdown (**Fig. 4d**).

ANME-2c FR_S4_sbin5 also harbor genes for complete MCR complex (McrABG), along with an archaeal Wood–Ljungdahl pathway for oxidizing methane to CO_2_ and HdrABC to recycle the CoM and CoB heterodisulfides (**Fig.4e and Fig.S12**). This denotes a conventional anaerobic oxidation of methane via MCR mechanism^64, 65^. The presence of C-P lyase in methanotrophs revealed the potential of methane production from utilizing methylphosphonate. Indeed, the metabolic product of PhnJ (alpha-D-ribose 1,2-cyclic phosphate 5-phosphate) was detected in our metabolome data (**Table S5**), indicating the actual occurrence of methylphosphonate-driven methane formation at cold seeps. All *phnJ* genes, encoding for the catalytic subunit^66^, from gene- and genome-level were used for phylogenic analyses to reveal the evolution of C-P layse. PhnJ protein sequences associated with Halobacteriota formed a monophyletic group, indicating a shared evolutionary ancestry (**Fig. S13**).

### Cold seep viruses regulate microbial phosphorus cycling via the PhoR-PhoB system and PhnCDE transporter

Apart from bacteria and archaea, we identified 134 phosphorus-cycling auxiliary metabolic genes (AMGs) encoded by 121 viral contigs (vContigs) from a cold seep gene catalog of 147 million non-redundant genes (**Table S7**). These vContigs were validated as viruses by containing viral hallmark genes, viral-like genes or having a high virus score (> 0.7) determined by genomad^67^. Genomic arrangements also supported their affiliations to viruses, with phosphorus-cycling AMGs flanked by viral hallmarks or viral-like genes on both sides (**Fig. 5a and Fig. S14**). Using geNomad^67^, up to 91.74% (n = 111) of the 121 vContigs were classified to the *Caudoviricetes* class, with a few assigned to classes of *Megaviricetes* (n = 1) and *Tokiviricetes* (n = 1, **Fig. 5b**). Due to the limited number of vContigs that could be classified to a specific viral family (**Table S8**), the family-level classifications were performed alternatively using PhaGCN^68^. A total of 49 vContigs could be further assigned to 10 families, with *Straboviridae* being the most prevalent (n = 19, **Fig. S15**).

**Figure 5.**
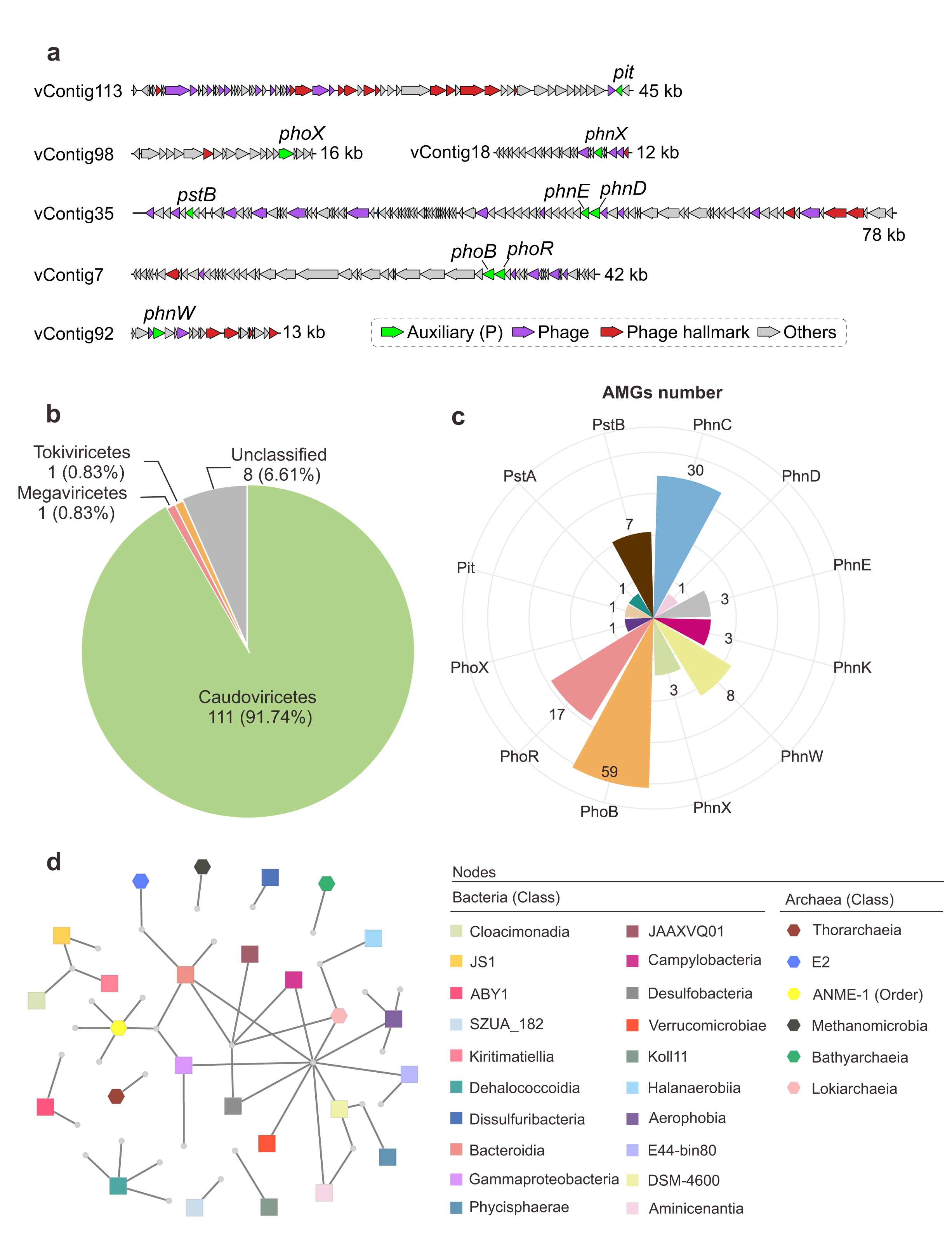
Phosphorus-cycling viruses at cold seeps. **a**, Genome organization diagrams of six representative vContigs identified in this study. Predicted open reading frames are colored according to the results of our phosphorus-cycling proteins annotation pipeline, DRAM-v and geNomad. **b**, The class-level classifications of the phosphorus-cycling vContigs according to geNomad. **c**, Numbers of the auxiliary metabolism genes (AMGs) involved in phosphorus cycling. **d**, Interaction networks of phosphorus-cycling vContigs and cold seep microorganisms. The circles and hexagons represent bacterial and archaeal host, respectively. Hosts are shown at the bottom of the panel and colored according to their taxonomy at class level (order level for ANME-1).

The 134 phosphorus-cycling AMGs spanned 12 gene types (**Fig. 5c**), categorized into three phosphorus metabolic processes: organic phosphorus mineralization (*phoX*, *phnW*, *phnX*, and *phnK*), phosphorus transportation (*phnC*, *phnD*, *phnE*, *pit*, *pstA*, and *pstB*), and phosphorus regulation (*phoR* and *phoB*). The identification of previously unrecognized AMGs^69^, i.e., *phnX*, *phoX*, and *phnK*, expanded the diversity of AMGs involved in phosphorus cycling. Remarkably, phosphorus regulation (*phoR*, n = 17; *phoB*, n = 59) was predominantly encoded by cold seep viruses **(Fig. 5c**). Moreover, one viral genome harbored a complete PhoB-PhoR regulatory system (**Fig. 5a**), a significant finding since previously only individual components of this system, either PhoR or PhoB, had been identified as AMGs^69, 70^. These finding allow us to propose that cold seep phages could regulate the transcription of phosphorus-cycling genes completely using their own regulators, rather than relying on the host’s regulatory system^71^. Phosphorus-cycling AMGs were also enriched in ATP-binding subunit (PhnC, n = 30; **Fig. 5c**) of the ABC-type phosphonate transport system (PhnCDE)^72^. One viral genome was observed to encode other components of this transporter (PhnDE, **Fig. 5a**). To date, only one provirus genome encoding PhnCDE was detected^73^, and the periplasmic high-affinity phosphate-binding gene *pstS* was the most commonly reported AMG related to phosphorus transport^74, 75^. The viral *phoR*, *phoB* and *phnC* were expressed at varying levels, with the highest reaching up to 4.8 TPM at Jiaolong cold seep, while the transcripts of other types of AMGs were not detected (**Table S9**). This enrichment of AMGs in phosphorus regulation and transport were consistent with the pattern observed in their microbial hosts, which had the highest abundance of these genes across global cold seeps (**Fig. 3a**). This uniformity suggests that cold seep viruses and their microbial hosts are likely under the same selective pressure (e.g., phosphate bioavailability) to retain these genes for phosphorus utilization, as seen with other nutrient cycling processes such as nitrogen and sulfur^76, 77^. Multiple lines of evidence suggest that, in phosphorus-limited oceanic water and various terrestrial ecosystems^69, 73, 78^, viruses augment their hosts’ phosphorus acquisition by encoding AMGs related to phosphatases like *phoD* and *phoN*. However, in phosphorus-rich cold seep environments, the dominance of AMGs for the PhoR-PhoB system and PhnCDE transporter implies that phosphorus regulation and transport are more crucial than metabolic processes for phosphate release.

To explore the potential viral impacts on phosphorus cycling at cold seeps, we investigated virus-host linkages, via iPHoP that aggregates six different methods for host predictions along with CRISPR spacer matching^79^. Approximate 26% (31/121) of the phosphorus-cycling vContigs identified could be associated with a total of 42 hosts (**Table S10**). These hosts encompassed a wide range of diversity, spanning 20 bacterial classes and 6 archaeal classes (order level for ANME-1; **Fig. 5d**). While it is established that known hosts for phosphorus-cycling viruses are predominantly bacteria^69^, our results revealed a broader spectrum of hosts in archaeal lineages. Specifically, we identified five distinct Caudoviricetes viruses infected ecologically important archaea ANME-1 (**Fig. 5d**), functioning as methanotrophs in many methane-rich ecosystems^7^. These viruses modulated the phosphorus metabolism of ANME-1 by containing *phoB*, *phnC*, and *phnK* (**Tables S7 and S10**). Of particular significance is the involvement of PhnK, a component of C-P lyase, in mediating methylphosphonate-driven methane formation^62^.

### Conclusion

Traditional homology-based annotations can only capture sequence space conserved to some degree, leaving a high number of genes unannotated and therefore ‘hidden’ from our sight. Here we show the utility of alignment-free PLMs to improve the functional prediction of phosphorus-cycling proteins within large-scale cold seep metagenomic datasets. The LucaPCycle model enabled the characterization of previously unrecognized alkaline phosphatase families featured by a streamlined core architecture domain and a unique C-terminal β-sheet domain. The discovery of these new enzyme families highlights the hidden diversity of phosphorus-cycling microbial communities and their overlooked ecological functions. The presence of diverse phosphoesters and phosphonates, along with their intermediate metabolites, and the high abundance and expression of phosphorus-cycling genes, all suggest that phosphorus cycling plays a significant yet often underappreciated role in the biogeochemical cycling within cold seep environments. Our study also expanded the taxonomic distribution of phosphorus-cycling genes among the Archaea domain. We highlight the unexpected roles of archaea in polyphosphate metabolism, mineral phosphorus solubilization, and both phosphonate substrate-specific and broad-specificity catabolism. Notably, ANME methanotrophs possess the genetic equipment for methylphosphonate-driven methane production via complete C-P lyase operons. Additionally, cold seep viruses regulate microbial phosphorus-cycling processes primarily through the PhoR-PhoB regulatory system and the PhnCDE transporter. This study underscores the critical need for PLM-based methodologies to uncover the hidden biological functions of these genes, enhancing our understanding of microbially-mediated phosphorus-cycling and its ecological significance in diverse ecosystems. Furthermore, the deep-sea sedimentary ecosystems, particularly beyond ocean water, represents a significant but often overlooked component in global phosphorus cycling.

## Methods

### Architecture of LucaPCycle

LucaPCycle is a transformer-based dual-channel model (**Fig. 1a**). The first stage (binary-classification) predicts if a protein sequence has a phosphorus-cycling function. In the second stage (31-classification), it classifies the positive predictions from the first stage into one of 31 specific types. These 31 phosphorus-cycling proteins participate in various phosphorus metabolisms, including phosphoester hydrolysis, phytate hydrolysis, broad-specificity and substrate-specific phosphonate catabolism, polyphosphate metabolism, mineral phosphorus solubilization, phosphonate transport, low- and high-affinity phosphate transport, and two-component systems (**Table S1**). LucaPCycle consists of five modules: Input, Tokenizer, Encoder, Pooler, and Output. Below are descriptions of each module:

1. Input: This module receives the amino acid sequence as input.
2. Tokenizer: This module performs tokenization of the input amino acid sequence at both character and word levels. Character-level tokenization slices protein sequences into amino acid characters. Word-level tokenization employs the Byte Pair Encoding (BPE) algorithm^80^ from the Subword-NMT tool (https://github.com/rsennrich/subword-nmt) to segment the input sequence into “words”, generating a word-level list.
3. Encoder: This module extracts features from the tokenized lists and includes two submodules. One submodule uses the ESM2-3B model^39^ to extract features from the character-level list, resulting in the representation matrix M1. This submodule captures universal features and internal structural and contextual information from the sequence. The second submodule based on the Transformer-Encoder ^40^ to derive features from the word-level list, producing the representation matrix M2.
4. Pooler: This module selects essential features and transforms the feature matrix into a vector through down-sampling. Since the Encoder extracts mixed features, especially general ones from ESM2, we use Value-Level Attention Pooling^81^ to assign learnable weights to each feature (Equations (1) and (2)).

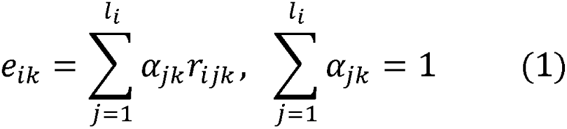

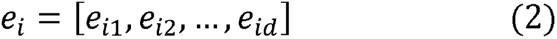

where, *i* is the sample index, *l_i_* is the length of the ;*i-th* sequence, *d* is the dimension of the embedding, *e_i_* is the pooled vector, and *k* ∈ {1,2,…,*d*}, *r_ijk_* is the value of row *j* and column *k* of the representation matrix of the *i-th* sequence.
5. Output: This module concatenates two pooled vectors into a classification vector. It uses the sigmoid-function for binary-classification to generate the positive probability and the softmax-function for 31-classification to output probabilities for each class.

### Dataset for LucaPCycle

We initially constructed a dataset with 214,193 positive samples of 31 phosphorus-cycling proteins and 4,129,874 negative samples. The positive samples are sourced from the PCycDB that represents a comprehensive and accurate database for analyzing phosphorus-cycling genes and microorganisms^30^. The number of sequences for each phosphorus-cycling protein detailed in **Fig. S1a**. To maintain a reasonable ratio of positive to negative samples (1:3.5), we used CD-HIT v4.8.1^82^ to reduce the redundancy of the negative samples with 50% amino acid identity, resulting in 751,331 non-redundant negative samples for model building. The negative samples encompass protein sequences involved in intracellular phosphorus metabolism (n = 36,661), nitrogen cycling (n = 8,447) and sulfur cycling (n = 63,525). To ensure a diverse set of negative samples, we also incorporated other unrelated functional proteins from KEGG (n = 63,606) and UniProt (n = 579,092), such as protein sequences from viral capsid, DNA replication, RNA transcription, carbon metabolism, signal transduction, and secondary metabolism. This diversity in negative samples aims to enhance the model’s annotation performance and generalization ability. The detailed numbers for each category of negative samples are shown in **Fig. S1b**.

### Model Building of LucaPCycle

For binary-classification model, all positive samples (n = 214,193) and non-redundant negative samples (n = 751,331) were randomly divided into training, validation, and testing datasets with a ratio of 0.99: 0.005: 0.005 (**Fig. S1c**). Regarding 31-classification model, the dataset consists of all positive samples without negatives. The sequences of each phosphorus-cycling protein were randomly divided into training, validation, and testing datasets with a ratio of 0.9: 0.05: 0.05.

The first channel of LucaPCycle applied ESM2-3B (https://github.com/facebookresearch/esm) for contextual representation. The second channel used a 4-layers with 8-heads Transformer-Encoder to characterize the raw sequence. The model training lasted for a maximum of 50 epochs based on the best F1-score in validation dataset for model finalization, with a learning rate of 1E-4, using a warm-up approach for updates. We employed a batch size of 8 with a gradient accumulation step of 8 for the binary-classification model and a batch size of 16 with a gradient accumulation step of 4 for the 31-classification model. The model was trained on a single Nvidia A100 GPU.

### Model evaluation and benchmarking

We evaluated binary-classification model and 31-classification model using six metrics based on validation and testing datasets, i.e., accuracy, precision, recall, F1-score, AUC, and PRAUC (**Fig. 1b**). To further benchmark LucaPCycle, we compared it against Diamond Blastp and KofamScan using a dataset of 1,463,634 true positive sequences (**Fig. 1c**). Precision and recall were calculated using equations (4) and (5).

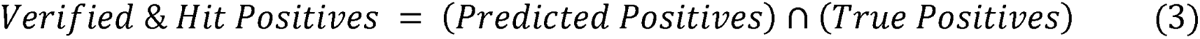

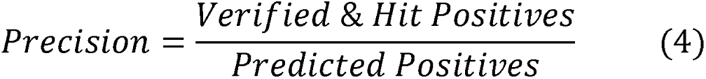

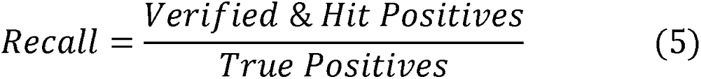

### Metagenomic and metatranscriptomic datasets and processing

The 165 metagenomes and 33 metatranscriptomes analyzed in this study were sourced from sediment samples (sediment depths: 0–68.55 mbsf; water depths 860–3,005 m) collected at 16 globally distributed cold seep sites (**Table S11**). These samples encompass five types of cold seeps, namely gas hydrates (n = 39), mud volcanoes (n = 7), asphalt volcanoes (n = 7), oil and gas seeps (n = 15) and methane seeps (n = 96).

Non-redundant gene and genome catalogs were constructed as described in our previous publication^83^. In brief, metagenomic raw reads were quality-controlled and assembled into contigs. The protein-coding sequences were predicted from contigs using Prodigal v2.6.3 (parameter: -meta)^84^, and then clustered at 95% amino acid identity using CD-HIT v4.8.1^82^. This process yielded a non-redundant gene catalog (n = 147,289,169). MAGs were recovered from contigs with the length longer than 1 kb using six binning tools, including MetaBAT2^85^, MaxBin2^86^, CONCOCT^87^, SemiBin^88^, VAMB^89^ and Rosella (https://github.com/rhysnewell/rosella). All produced MAGs were dereplicated at 95% average nucleotide identity using dRep v3.4.0 (parameters: -comp 50 -con 10)^90^, resulting in 1878 representative MAGs. Completeness and contamination of MAGs were evaluated using CheckM2 v1.0.1^91^. Functional annotation of MAGs was carried out by searching against KEGG, Pfam, MEROPS and dbCAN databases using DRAM v1.3.5^92^.

With regard to 33 metatranscriptomes, raw reads were quality filtered using the Read_qc module within metaWRAP v1.3.2^93^. The removal of ribosomal RNAs was conducted using sortmeRNA v2.1 with default parameters^94^.

### Identification of phosphorus-cycling proteins based on LucaPCycle and two homology-based methods

We used the trained LucaPCycle to predict 31 phosphorus-cycling proteins from the protein sequences of non-redundant gene catalogue and MAGs. Detailed predictions for each protein are shown in **Fig. S1d**. Additionally, the same datasets were also functionally annotated using two conventional methods, blast- and HMM-based approaches. For the blast-based approach, we used Diamond Blastp program v2.0.15^95^, with an identity ≥ 30.0% and hit length ≥ 25 amino acids. For the HMM-based approach, we used pre-computed HMM profiles of KEGG ortholog via KofamScan v1.3.0^31^. The significant hits related to 31 phosphorus-cycling genes were selected according to KofamScan’s adaptive score threshold and an e-value threshold of 1E-5.

To ensure more precise and reliable results, all protein sequences identified by LucaPCycle, Diamond Blastp, and KofamScan were merged, followed by validation through three distinct methods: domain analysis, DeepFRI v1.0.0 (Deep Functional Residue Identification)^41^, and CLEAN v1.0.1 (Contrastive Learning-Enabled Enzyme Annotation)^42^. With regard to domain analysis, protein sequences were queried against full ECOD domain database (v.20231128)^96^ using DIAMOND blastp program v2.0.15^95^ with an e-value threshold of 1E-2. ECOD is a hierarchical domain database that describes the evolutionary relationships between pairs of protein domains. A sequence was considered as bona fide phosphorus-cycling protein if supported by at least one of these validation processes. The cross-checked results were used for benchmark test.

To reveal phylogenetic diversity, clustering was conducted on the validated protein sequences using CD-HIT v4.8.1 (parameters: 70% sequence identity, 80% alignment coverage)^82^ to select representative ones. At this step, all singleton clusters were removed, resulting in 321,789 non-singleton protein families associated with phosphorus-cycling proteins.

### Geochemical and metabolomics analyses

The porewater samples from Qiongdongnan cold seep were used for the following geochemical analyses. Total alkalinity (TA) was analyzed onboard by direct titration with an HCl standard solution. The pH of the porewater was monitored using a calibrated pH meter. Phosphate (PO_4_^3-^) concentrations were determined photometrically using a UV-Vis spectrophotometer (Hitachi U5100, Hitachi Limited, Japan). Other cation-anion concentrations (SO ^2-^, K^+^, Na^+^, Mg^2+^ and Ca^2+^) were measured using an ion chromatography system (ICS-1100, Thermo, United States) with conductance detector. The total organic carbon (TOC) and total nitrogen (TN) of bulk sediment samples were quantified on an elemental analyzer-isotope ratio mass spectrometer (Vario ISOPOTE Cube-Isoprime, Elementar, Germany).

The metabolites were extracted from 55 sediment samples collected from the Qiongdongnan and Shenhu cold seeps following previously reported methods^97^. High-resolution LC-MS/MS analysis was performed on a Waters Acquity I-Class PLUS ultra-high performance liquid tandem with a Waters Xevo G2-XS QTOF high-resolution mass spectrometer. Data collection was performed in MSe mode and processed using Progenesis QI software, utilizing the METLIN database and Biomark’s proprietary library for metabolite identification.

### Taxonomy assignment of Bacteria and Archaea

To obtain the taxonomic profiles for microbial communities driving the process of phosphorus cycling, all sequences from non-singleton phosphorus-cycling protein families were taxonomically assigned using the MMseqs2 taxonomy tool v13.45111 (parameter: --tax-lineage 1)^98^. This involved performing six-frame translation searches against the GTDB R207 database and assigning each protein sequence to the lowest common ancestor of the best hits for each frame.

The taxonomic classification of MAGs encoding phosphorus-cycling proteins was conducted using GTDB-Tk toolkit v2.1.1^99^ with default parameters against the GTDB R207database. The assignment of MAG FR_S4_sbin harboring both McrA and PhnJ were further confirmed by the visual inspection of taxonomic trees. Briefly, the concatenated multiple sequence alignment of the MAG FR_S4_sbin and reference ANME genomes accessed from NCBI GenBank were produced based on 43 conserved single-copy genes extracted by CheckM v1.0.12^100^. The maximum likelihood phylogenomic tree was generated using RAxML v8.2.12^101^ with PROTCATLG model and 1000 bootstrap replicates.

### Abundance calculations

The gene abundances of non-singleton phosphorus-cycling protein families across 165 metagenomes were quantified using Salmon v.1.9.0 in mapping-based mode (parameters: -validateMappings -meta)^102^. GPM (genes per million) values were used as a proxy for gene abundance. Similarly, transcript abundances of non-singleton phosphorus-cycling protein families were determined by mapping clean reads from 33 metatranscriptomes to the non-redundant gene catalog using Salmon v.1.9.0 with the aforementioned parameters^102^. Transcript abundances were calculated as TPM (transcripts per million). It is worth noting that all sequences from non-singleton phosphorus-cycling protein families were considered in these calculations.

### Protein structure alignments, domain classification and molecular docking

Three-dimensional structures of identified phosphorus-cycling proteins were predicted using the deep-learning artificial intelligence system, via AlphaFold3 web server (May 2024 to June 2024)^103^. The confidence in the accuracy of the predicted structures was measured by the predicted template modeling (pTM) score. All remote ALP structures exhibited pTM score above 0.5 and 76% of the structures achieving very high confidence scores (pTM > 0.9; **Table S12**). A pTM score above 0.5 means the overall predicted fold for the protein might be similar to the true structure. For PhnGHIJ complex, the interface predicted template modeling (ipTM) score was used to measure the accuracy of the predicted relative positions of the subunits within this complex.

The paired structural alignments were carried out using Foldseek v8.ef4e960 ^33^ in easy-search mode to align the structures of remote alkaline phosphatases against a library of reference structures drawn from PDB database and manually curated AlphaFold models (**Table S13**). The domains of remote ALPs were analyzed with FoldSeek v8.ef4e960^33^ by comparing to domains defined in the ECOD domain databases (v.20231128)^96^.

Ledock (v1.0; https://www.lephar.com/) was used to predict the binding poses of 89 phosphoester compounds on remote ALP (RMSD: 1.0, number of binding conformations: 20). The binding box of each predicted protein was performed via the DockingPie1 plugin within PyMOL (docking program: Autodock Vina)^104^. All of the structures were visualized and exported as images using PyMOL.

### Phylogenetic analysis

To facilitate a clear and intuitive phylogenetic tree, we used remote ALPs representative sequences from non-singleton phosphorus-cycling protein families. For sequence-based tree, the sequences of remote ALPs were aligned with corresponding reference sequences using MUSCLE v3.8.1551 (default settings)^105^. The alignments were subjected to phylogenetic tree reconstruction using FastTree v2.1.11 with JC+CAT models^106^. Structures of remote ALPs derived from AlphaFold 3 modeling, along with reference structures of well-characterized ALPs were subjected to structural tree building using Foldtree (https://github.com/DessimozLab/fold_tree), based on a local structural alphabet. All trees were visualized using iTOL v6^107^.

### Identification of viral genomes-containing phosphorus-cycling genes

The assembled contigs with length longer than 10 kb were screened by geNomad pipeline v1.3.3 (parameter: genomad end-to-end)^67^ for viral prediction. The completeness and contamination of identified viral genomes were determined by CheckV v1.01 (database v1.5)^108^. Only viral genome-containing phosphorus-cycling genes were included for downstream analyzes. The taxonomy each of phosphorus-cycling viral genome was assigned by geNomad v1.3.3^67^. However, only a few viral genomes could be assigned to specific viral families or genera. Thus, the finer classifications of phosphorus-cycling viral genomes were alternatively performed by PhaGCN v2.0 (cut-off score > 0.5)^68^. Phage genes and hallmark genes were identified by DRAM-v v1.3.5^92^ and geNomad v1.3.3^67^. The putative virus-host linkages were predicted using iPHoP v1.3.3^79^ and CRISPR spacer matching. For the iPHoP, both default database and custom MAGs from our 165 metagenomes were used to maximize host prediction. For CRISPR spacer matching, the CRISPR arrays were identified using CRISPRCasFinder v4.2.21^109^ from our custom MAGs. Local alignments of extracted spacers with lengths greater than 25 bp against viral genomes were searched using BLASTn. Only BLAST matches with 100% alignment coverage and at most two mismatches were considered as high-confidence matches.

### Statistical analysis

All statistical analyses were performed in R (v4.1.0). The normality and variance homogeneity of data were evaluated using Shapiro-Wilk test and Levene’s test, respectively. The Kruskal-Wallis rank-sum test was employed to compare gene abundance among different phosphorus metabolic processes. Correlation network analyses of environmental factors and phosphorus-cycling gene abundance were based on Spearman’s correlation. Pearson’s correlation and linear regression were performed to assess the relationship between PO_4_^3−^ concentrations and sediment depth. The alpha diversity (Chao1 index) of phosphorus-cycling microbial communities was calculated using the ‘vegan’ package. Variations in alpha diversity across different depths and cold seep types were evaluated using the Wilcoxon rank sum test and the Kruskal-Wallis rank sum test, respectively. The bata diversity of phosphorus-cycling microbial communities was estimated using the partial least squares discrimination analysis (PLS-DA) with the ‘mixOmics’ package. PERMANOVA (Permutational Multivariate Analysis of Variance; permutations = 999) tests were used to calculate the statistical differences across different depths and types of cold seeps. Both alpha and beta diversity indices were calculated based on functional profiles.

## Acknowledgements

The work was supported by the National Science Foundation of China (No. 32300100 and No.92351304), the Natural Science Foundation of Fujian Province (No.2023J06042), Scientific Research Foundation of Third Institute of Oceanography, MNR (No. 2022025 and No. 2023022), State Key Laboratory of Marine Geology, Tongji University (No. MGK202303), and China Postdoctoral Science Foundation (No. 2022M723709). We thank Jiaxue Peng and Yingchun Han for their assistance in metabolomics and metagenomic analyses, respectively, and Fabai Wu, Yu Hu, Muhe Diao, Maxim Rubin-Blum and Lingwei Ruan for helpful discussions.

## Author contributions

Conceptualization, X.D., and C.Z.; Methodology, C.Z., Y.H., Z.-R.L., J.W., T.C., J.L., and X.X.; Investigation, C.Z., X.D., Y.H., and Z.-R.L.; Computational Resources, Y.H., and Z.-R.L.; Writing – Original Draft, C.Z., Y.H., X.D., and Z.-R.L.; Writing – Review and Editing, X.D., C.Z., Y.H., Z.-R.L., M.H., and F.B.; Funding Acquisition, X.D., and C.Z.; Supervision, X.D., and Z.-R.L.

## Code & Data availability

The original source code for LucaPCycle is available at <u>LucaPCycle GitHub</u> or <u>LucaPCycle FTP</u>. The raw data and the processed dataset used for model building as well as the results from three functional annotation tools (i.e., LucaPCycle, Blastp and KofamScan) can be accessed at <u>LucaPCycle Data FTP</u>. Additional data produced in this study including representatives from non-singleton phosphorus-cycling protein families along with phylogenetic tree files based on amino acid sequences and protein structures of ALPs are available at <u>Tree-Families FTP</u>. The non-redundant gene and MAGs catalogs derived from 165 cold seep metagenomes can be found in Figshare (https://doi.org/10.6084/m9.figshare.22568107). See README on the <u>LucaPCycle</u> <u>FTP</u> for a detailed description of the code and data.

