## Supplementary Figures for "Illuminating microbial phosphorus cycling in deep-sea cold seep sediments using protein language models"

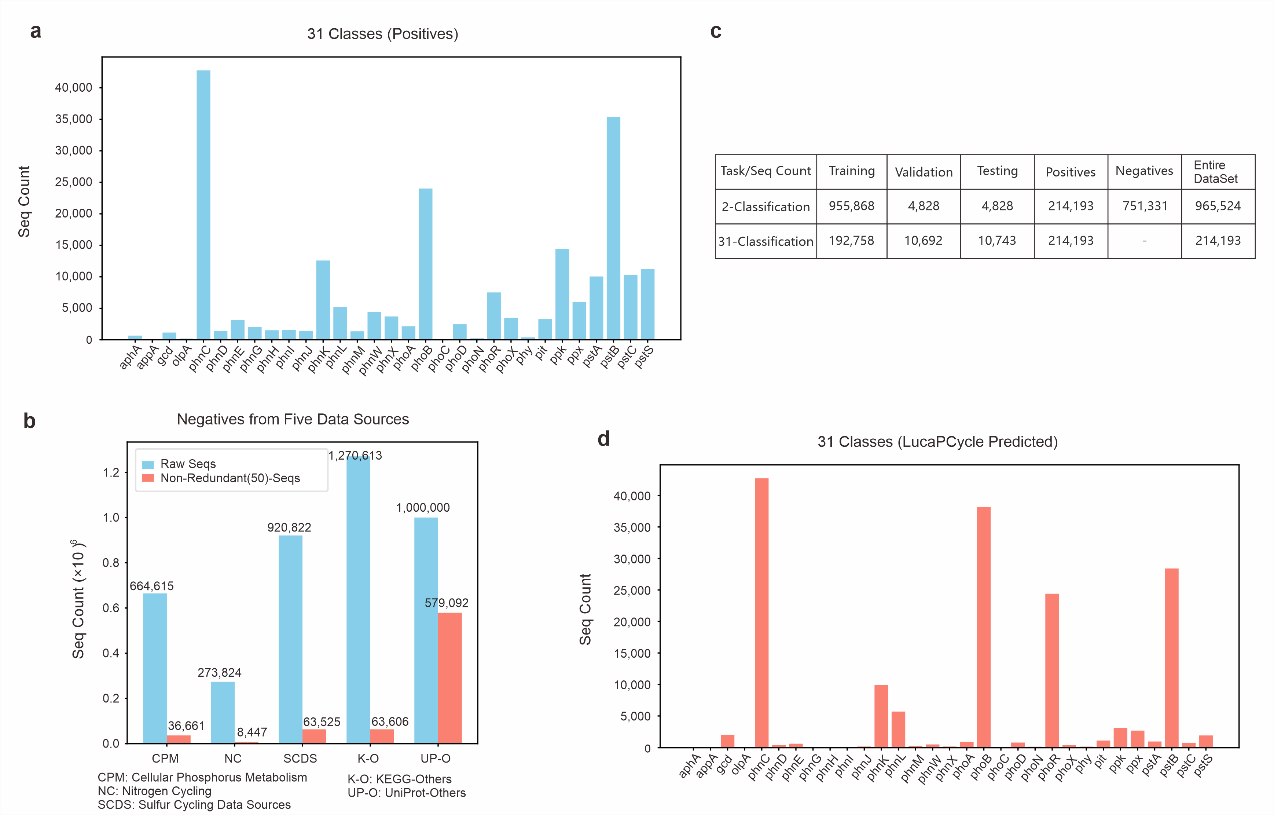

**Figure S1. Overview of model building datasets, training datasets, and predictions by LucaPCycle.** **a**, The number of sequences for each phosphorus-cycling proteins in a total of 214,193 positive samples. **b**, The number of negative sequences from different data sources. **c**, The number of samples divided into training, verification, and testing datasets for binary- and 31-classification models. **d**, The predicted numbers for each phosphorus-cycling protein by LucaPCycle.

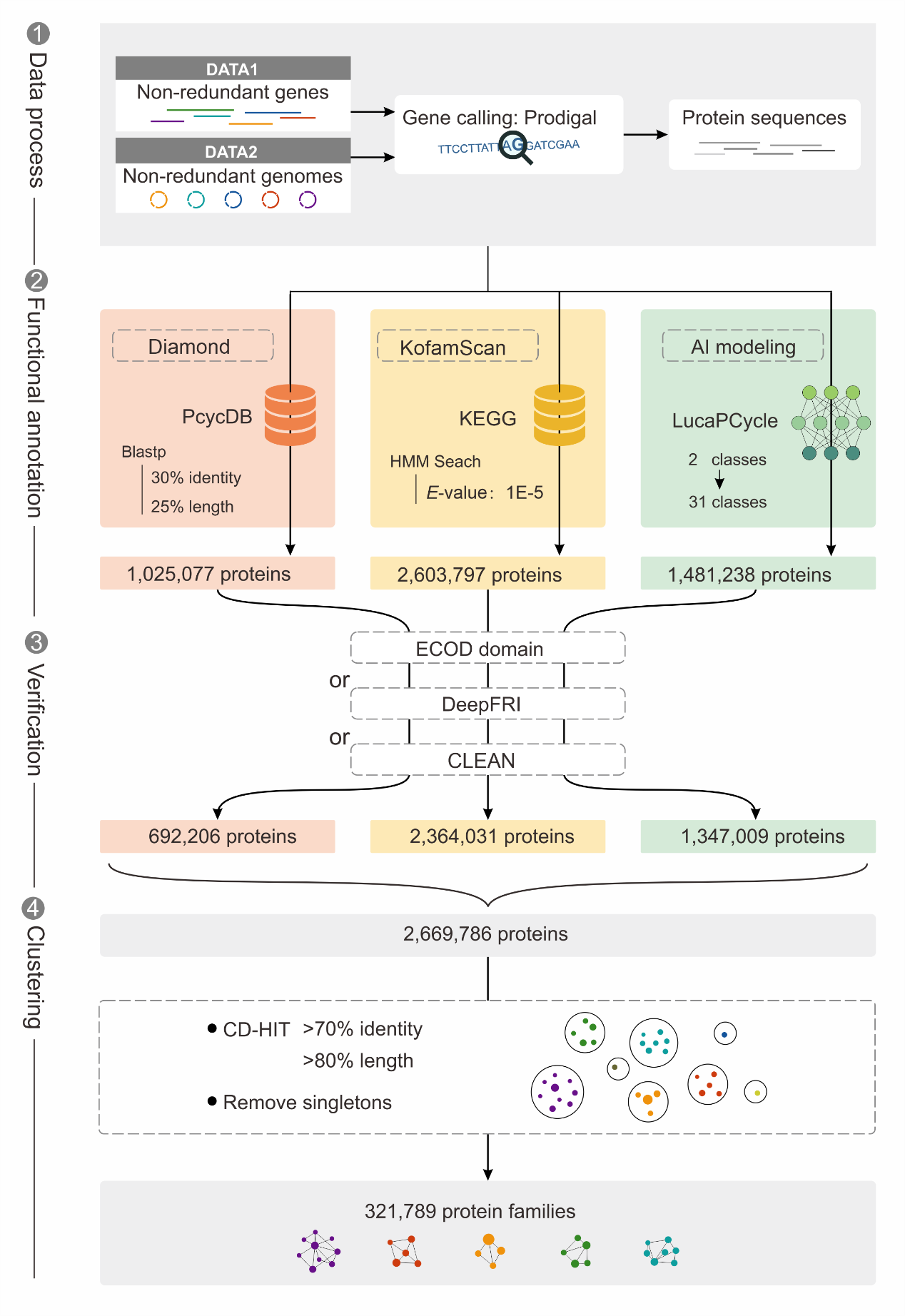

**Figure S2. Schematic diagram of LucaPCycle modelling and two sequence homology-based methods for phosphorus-cycling proteins.** Sequence-based searches include Diamond Blastp and KofamScan.

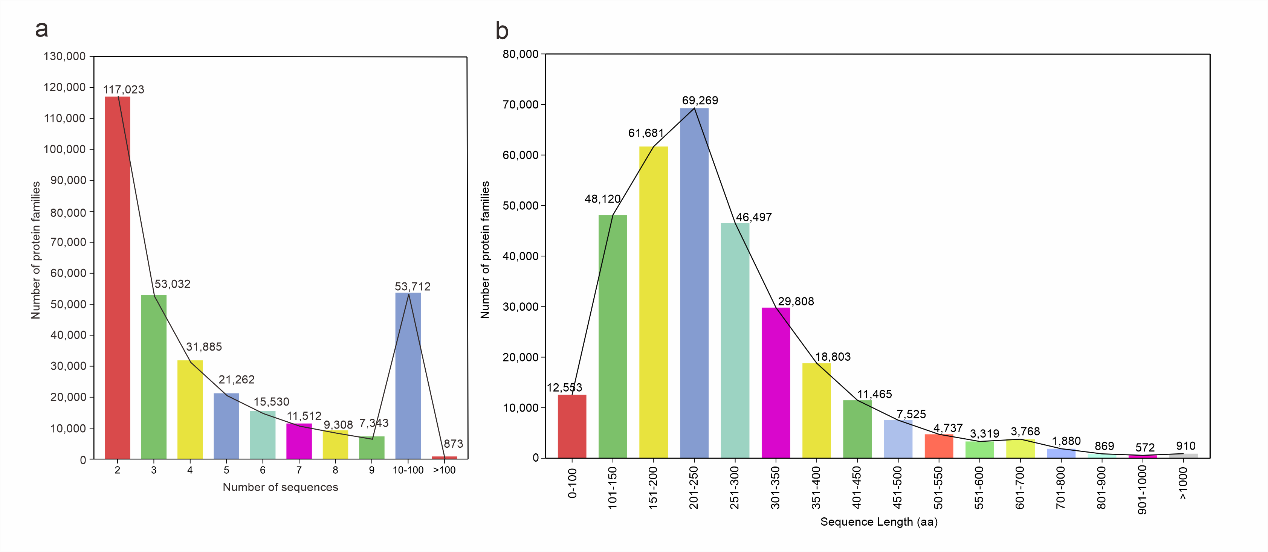

**Figure S3. The statistics of non-singleton phosphorus-cycling protein families.** **a**, Bar chart visualization of cluster components per cluster for the number of protein sequences in non-singleton phosphorus-cycling protein families. **b**, Bar chart distribution of the length of protein sequences in non-singleton phosphorus-cycling protein families. The horizontal axis shows the sequence length in amino acids, while the vertical axis shows the number of sequences in each length range.

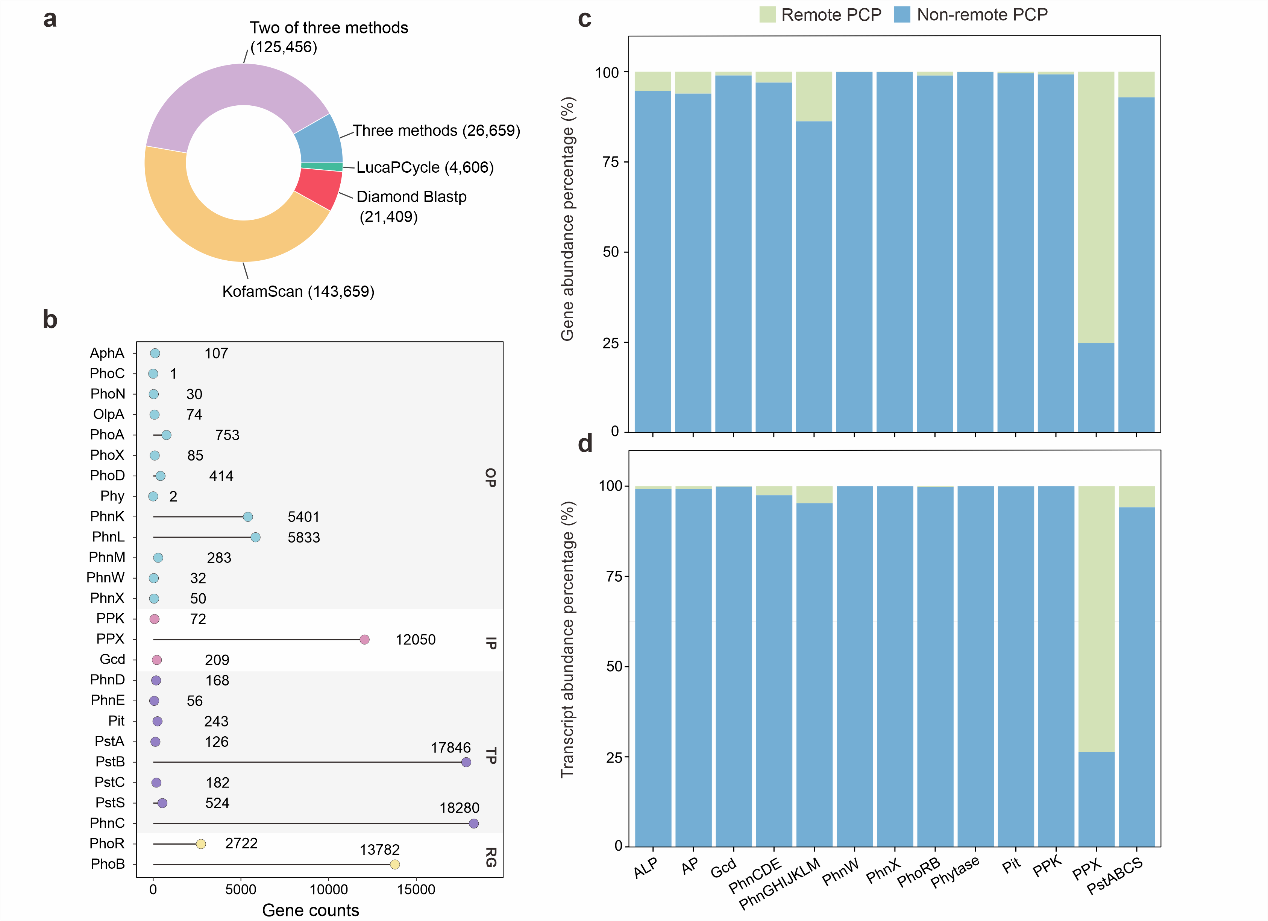

**Figure S4. Remote phosphorus-cycling proteins identified by LucaPCycle. a**, Distribution of protein families identified by LucaPCycle, Diamond Blastp and KofamScan. **b**, Numbers of the phosphorus-cycling genes with remote homology. OP: Organic phosphorus mineralization; IP: Inorganic phosphorus solubilization; TP: Transporters; RG: Regulatory genes. **c**, Percentage of remote phosphorus-cycling gene abundance. **d**, Percentage of transcript abundance of remote phosphorus-cycling gene. PCP: phosphorus-cycling proteins. ALP: alkaline phosphatase including PhoA, PhoD and PhoX. AP: acid phosphatase including AphA, OlpA, PhoN and PhoC.

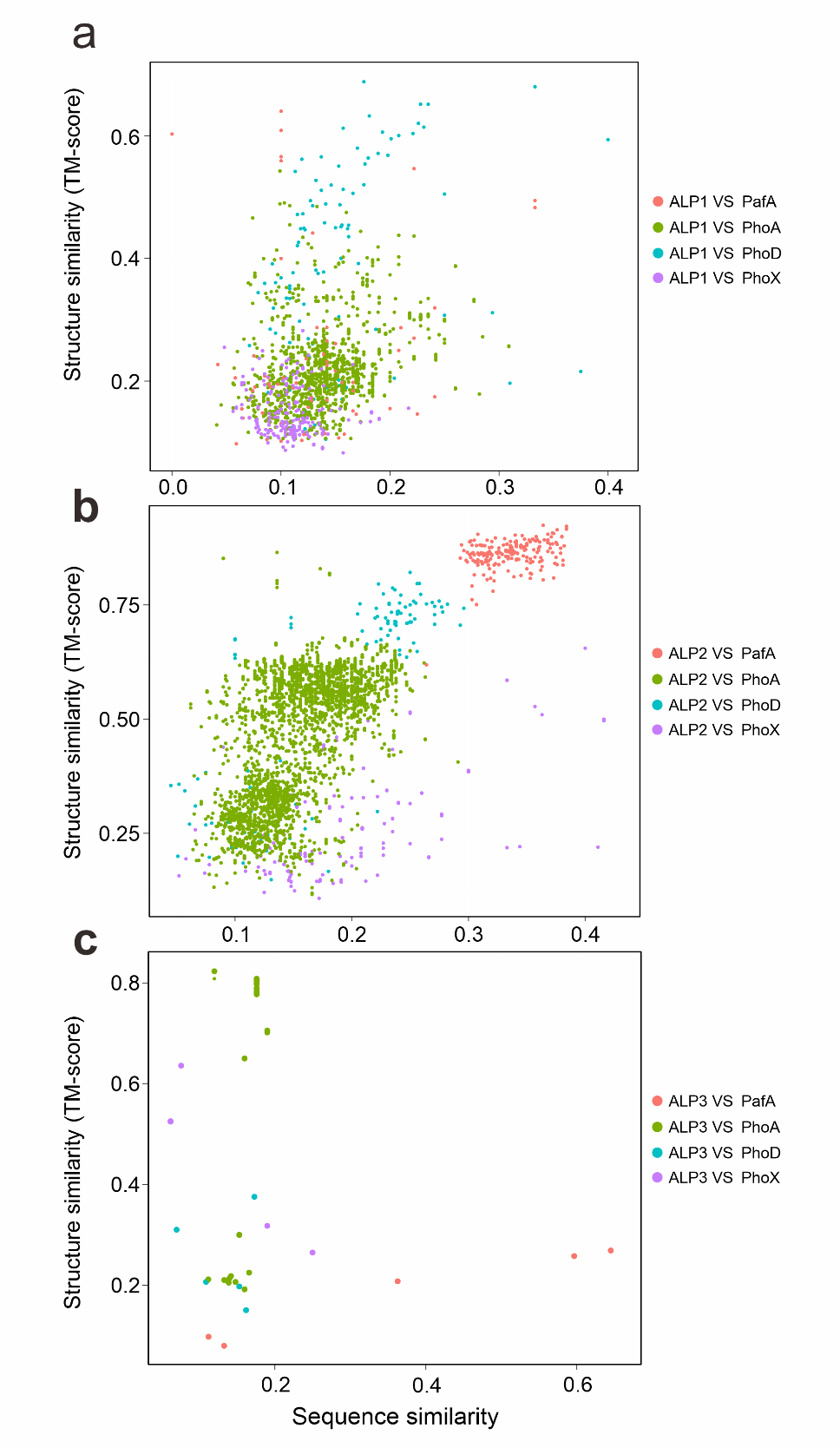

**Figure S5. Sequence similarity and structural similarity of ALP1 (a), ALP2 (b) and ALP3 (c) against three classical ALPs alongside the divergent PafA.** ALP: alkaline phosphatases including PhoA, PhoD and PhoX.

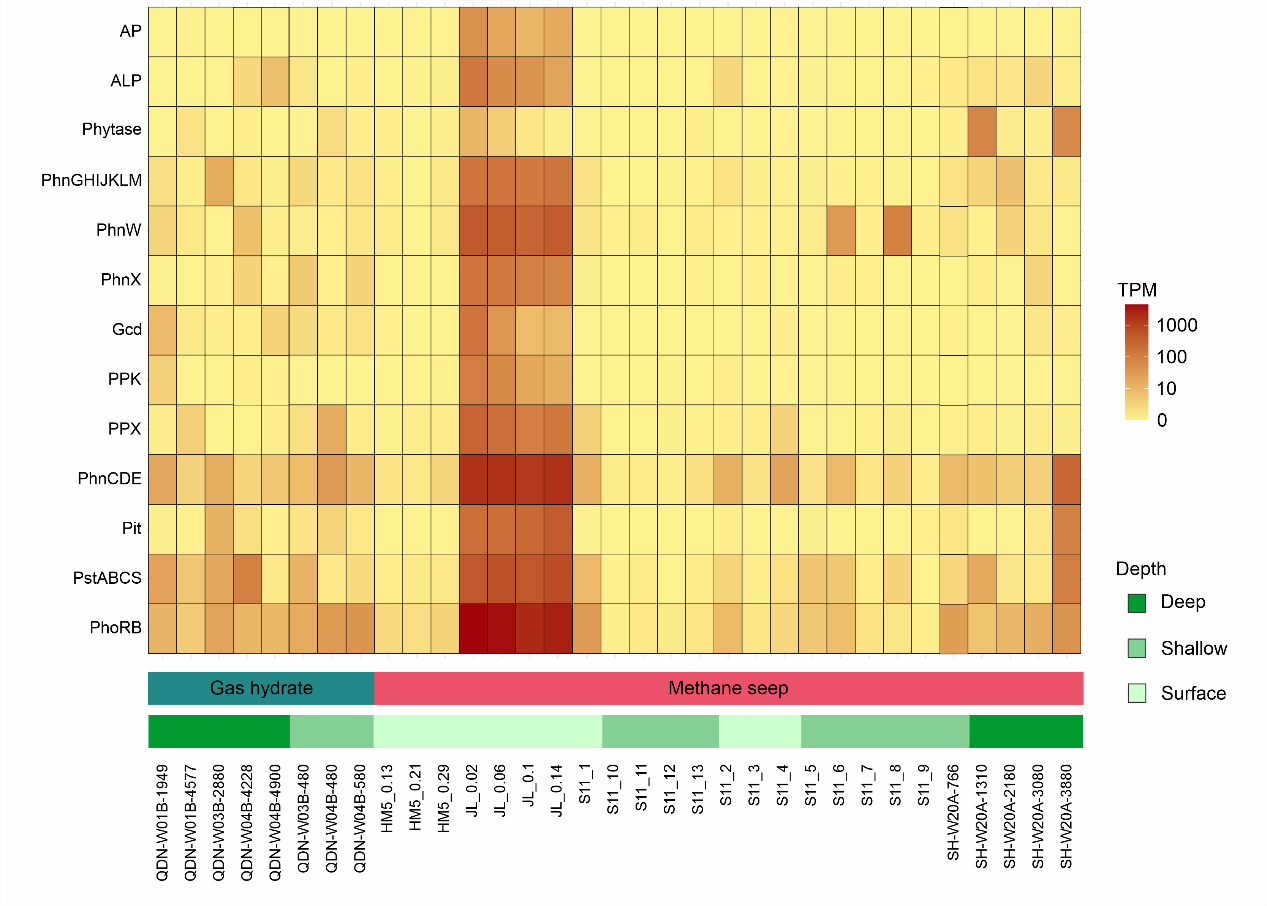

**Figure S6. The transcript abundances of phosphorus-cycling genes across the 33 cold seep metatranscriptomes**. Transcripts abundances are indicated as TPM (transcripts per million). Surface, < 1 mbsf; Shallow, 1 to 10 mbsf; Deep, > 10 mbsf. ALP: alkaline phosphatase including PhoA, PhoD and PhoX. AP: acid phosphatase including AphA, OlpA, PhoN and PhoC.

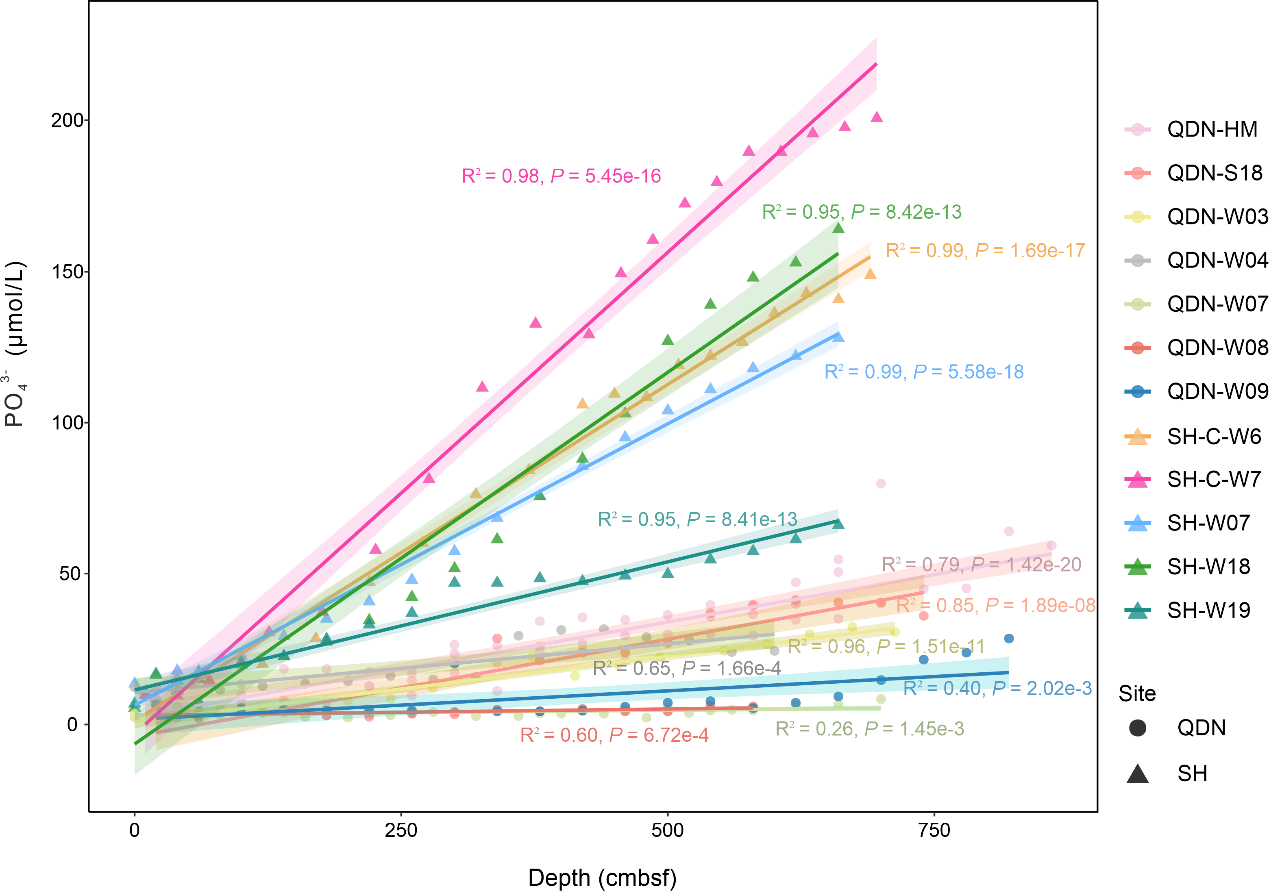

**Figure S7. Depth profiles of phosphate (PO_4_^3-^) concentration in 12 sediment cores collected from Qiongdongnan (QDN) and Shenhu (SH) cold seeps**. Each point represents the phosphate (PO_4_^3-^) concentration for a porewater sample, accompanied by a linear regression line and *R*^2^ value specific to each sediment, depicted in the corresponding color.

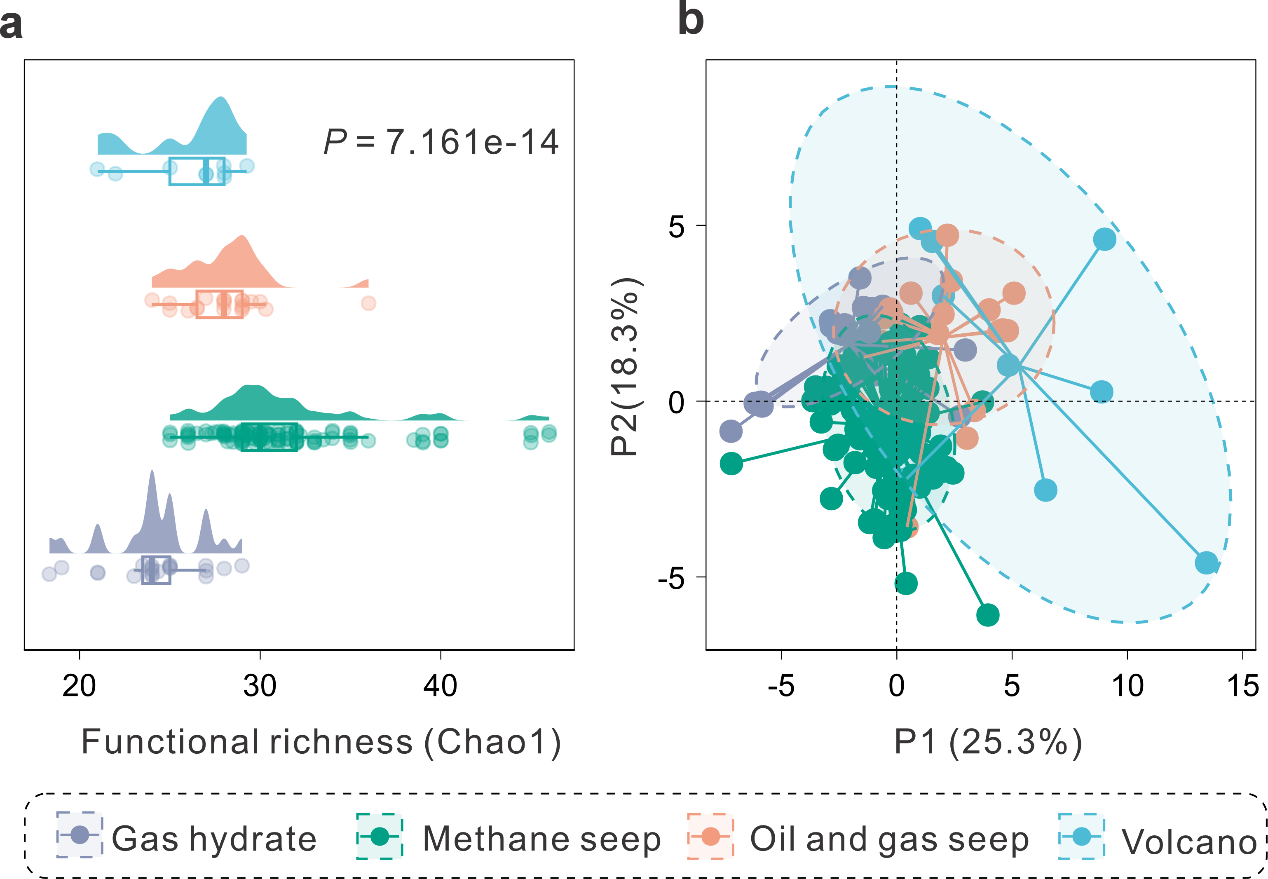

**Figure S8. The comparison of phosphorus-cycling proteins among cold seep types. a**, The functional richness of phosphorus-cycling genes across diverse cold seep types. **b**, The partial least squares discrimination analysis (PLS-DA) plot depicts the differences in phosphorus-cycling microbial communities among diverse cold seep types. Analyses of similarity among different types of cold seep were examined using a 999-permutation PERMANOVA test.

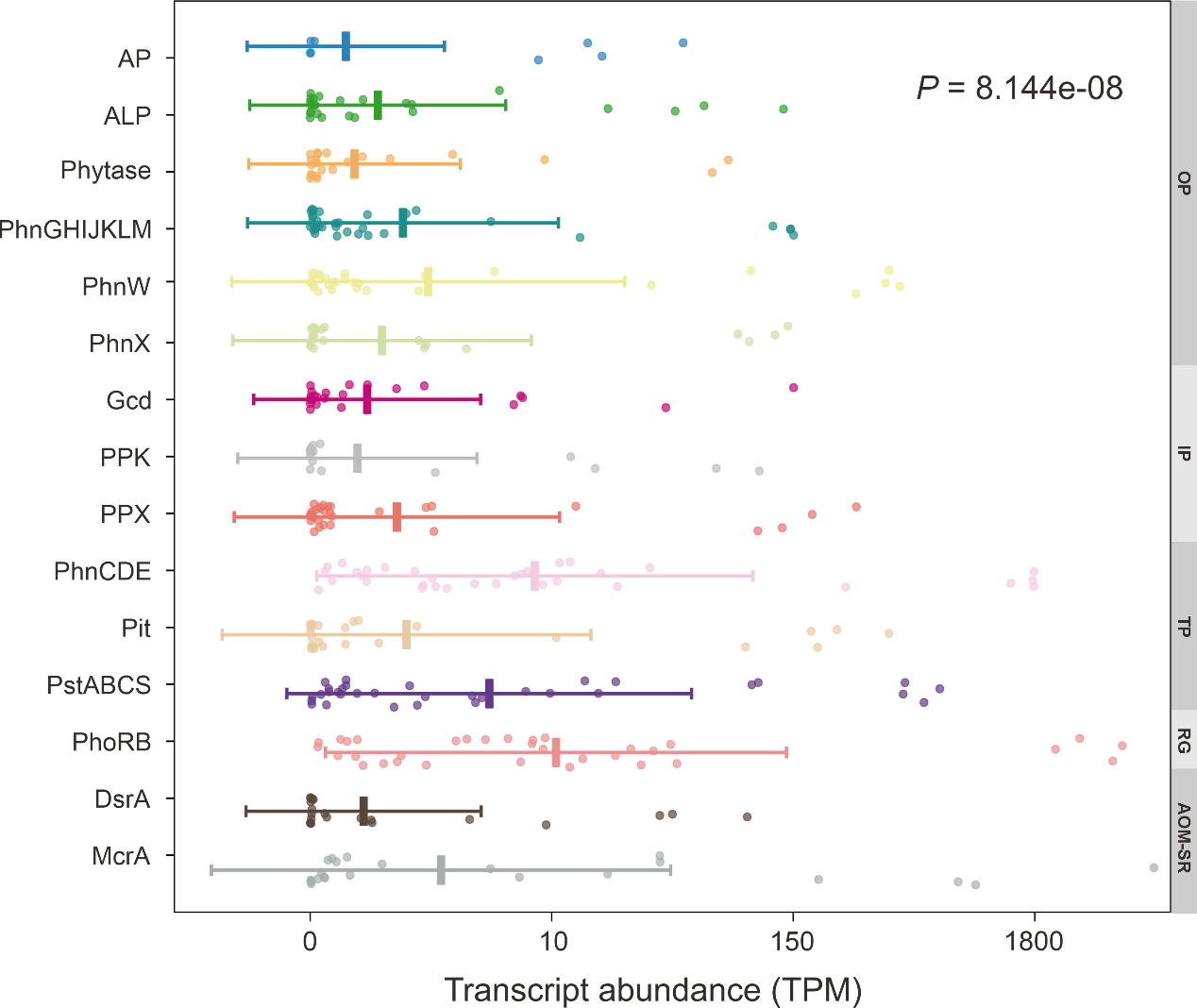

**Figure S9. Transcript abundances of phosphorus-cycling genes from four distinct metabolic processes, as compared to *mcrA* and *dsrA***. Transcripts abundances are indicated as TPM (transcripts per million). OP: Organic phosphorus mineralization; IP: Inorganic phosphorus solubilization; TP: Transporters; RG: Regulatory genes; AOM-SR: The anaerobic oxidation of methane (AOM) coupled to sulfate reduction (SR). ALP: alkaline phosphatase including PhoA, PhoD and PhoX. AP: acid phosphatase including AphA, OlpA, PhoN and PhoC. *P* values of differences among different genes were calculated using Kruskal-Wallis rank-sum tests.

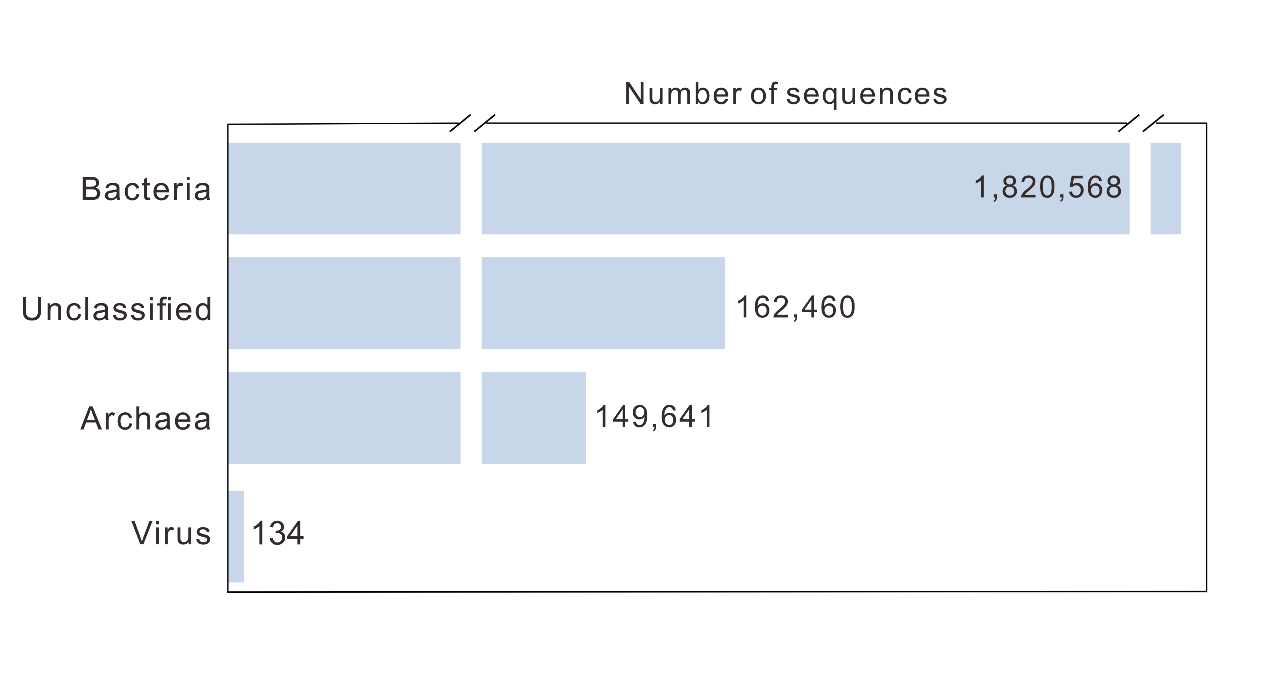

**Figure S10. Domain-level taxonomic distribution of non-singleton phosphorus-cycling protein families.** The total size of each classification category is represented by horizontal bar charts, with numbers labeled on the right side of the bars.

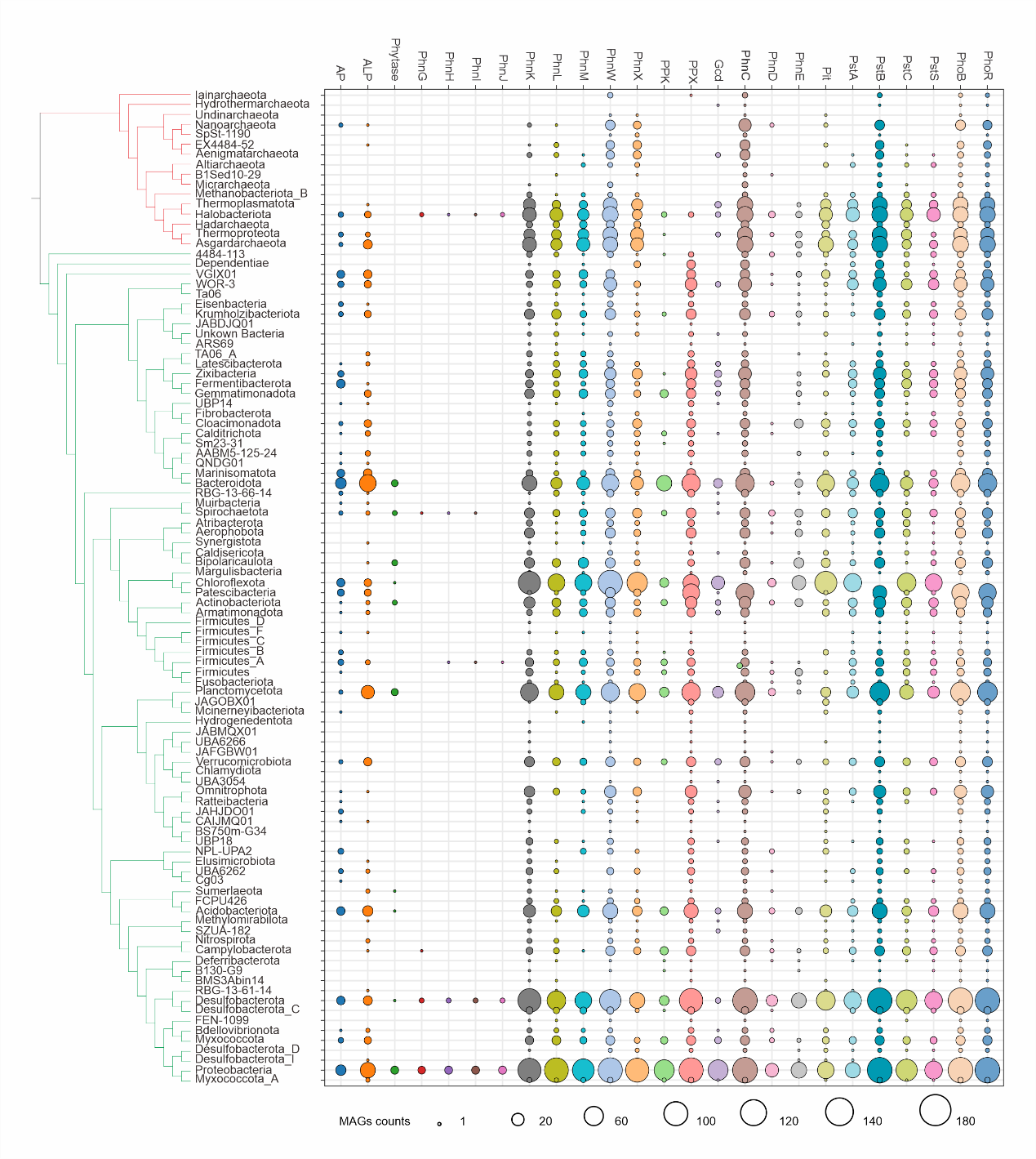

**Figure S11. Phylogenetic distribution of phosphorus-cycling genes across Archaea and Bacteria domain**. The right phylogenomic trees showing major lineages of Archaea and Bacteria. The tree was reconstructed from concatenate alignment of 43 conserved single-copy genes extracted by CheckM v1.0.12. The right bubble plot showing the number of phosphorus-cycling genomes within each phylogenetic cluster.

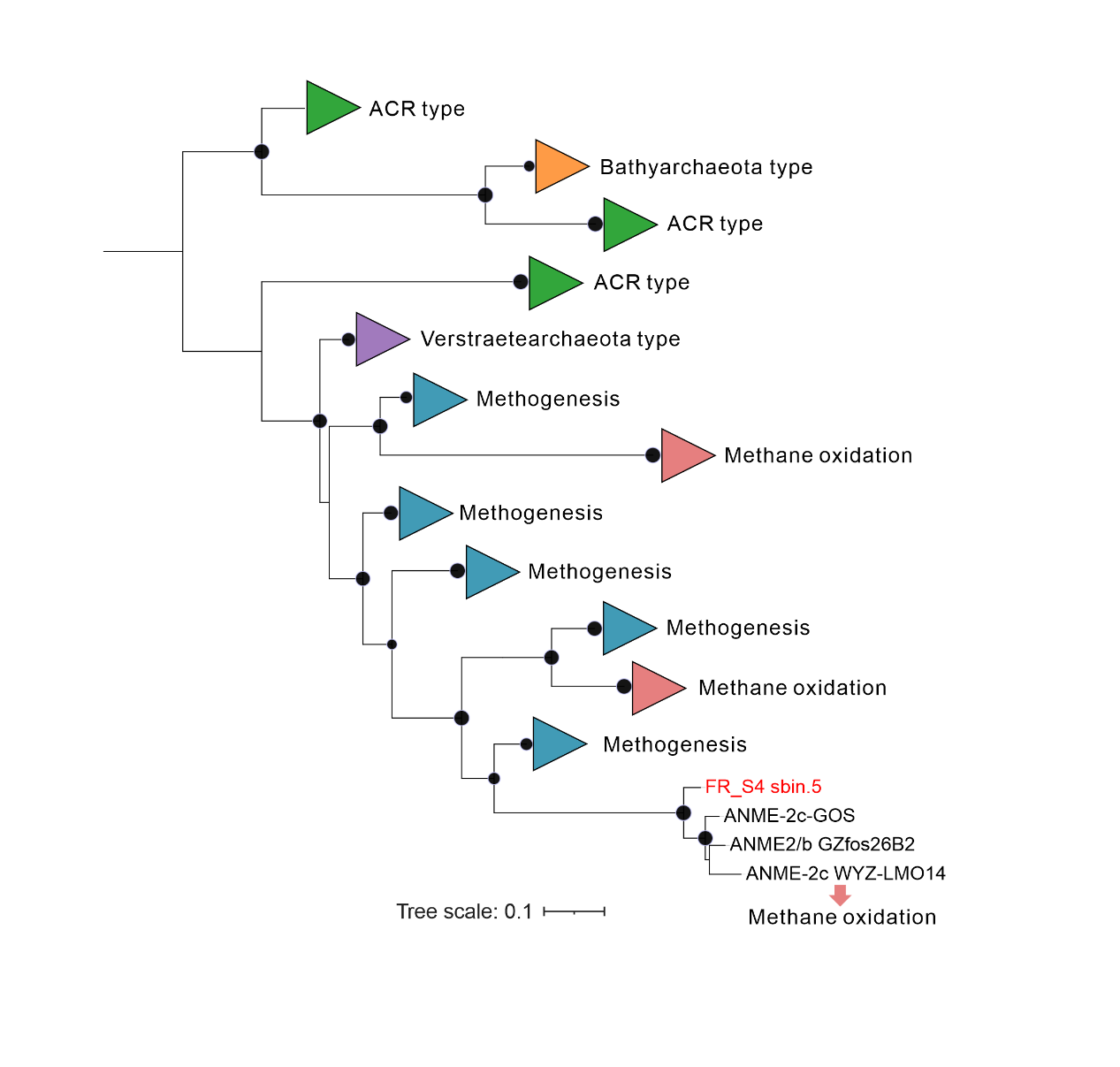

**Figure S12. Phylogenetic tree of McrA protein sequences recovered MAG FR_S4_sbin5**. Bootstrap value ≥70 are shown. Scale bar indicates amino acid substitutions per site.

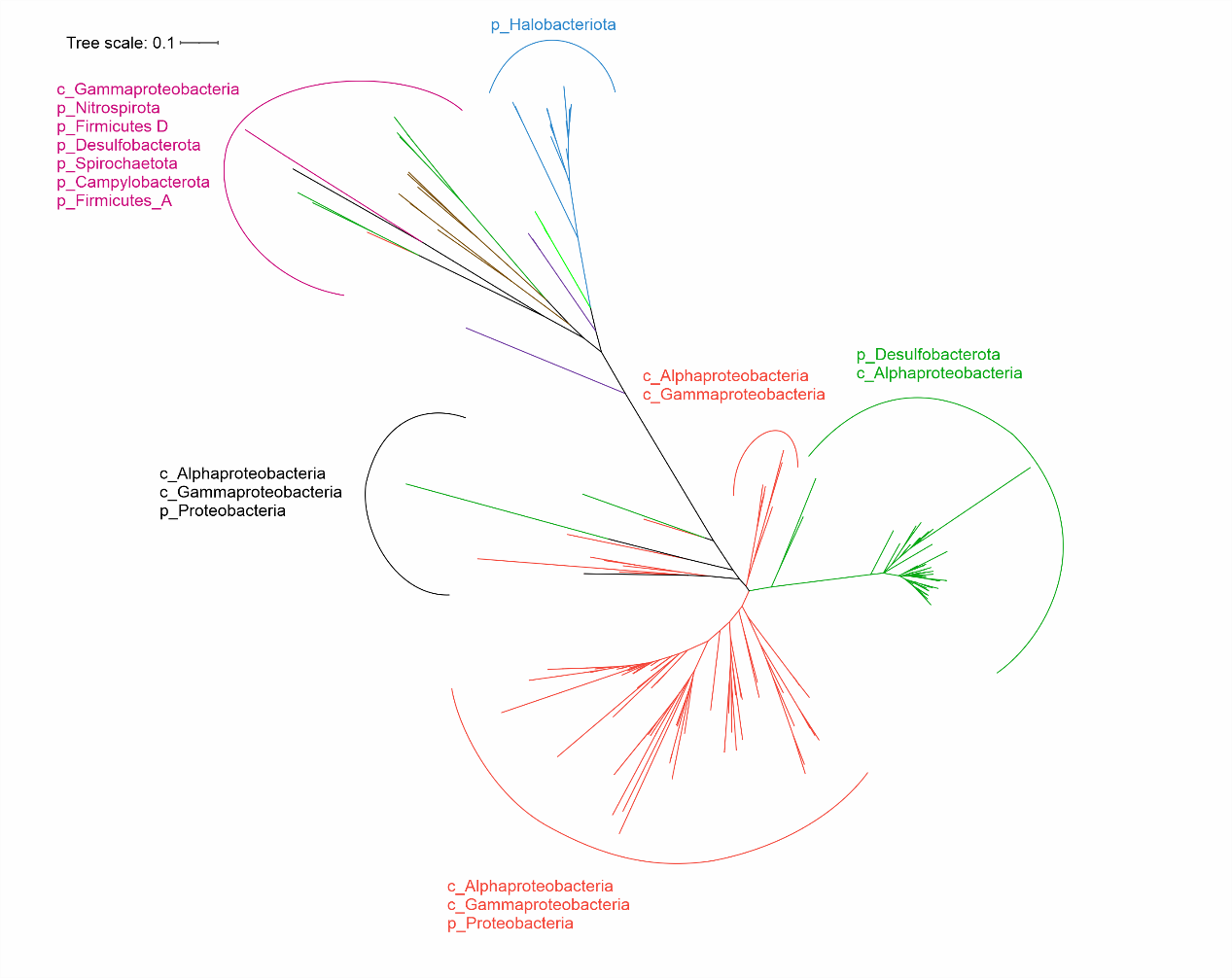

**Figure S13. Phylogenetic tree of PhnJ protein sequences.** Scale bar indicates the mean number of substitutions per site. Bootstrap values over 70% were shown. The GTDB taxonomy is indicated by the color code alongside the tree**.**

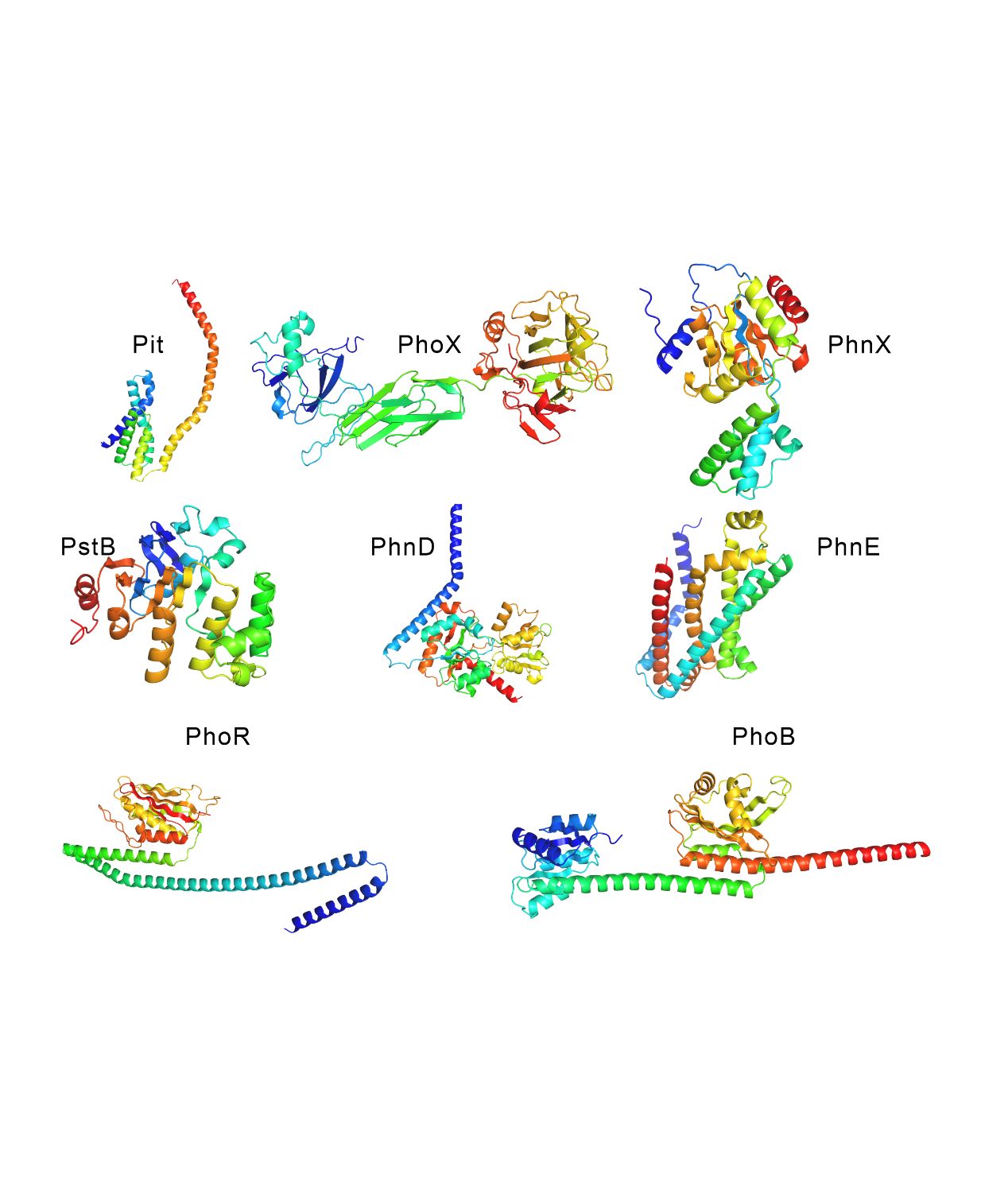

**Figure S14. Predicted protein structures of P-solubilizing AMGs**. The protein structures are predicted by AlphaFold 3 and represented in a rainbow spectrum.

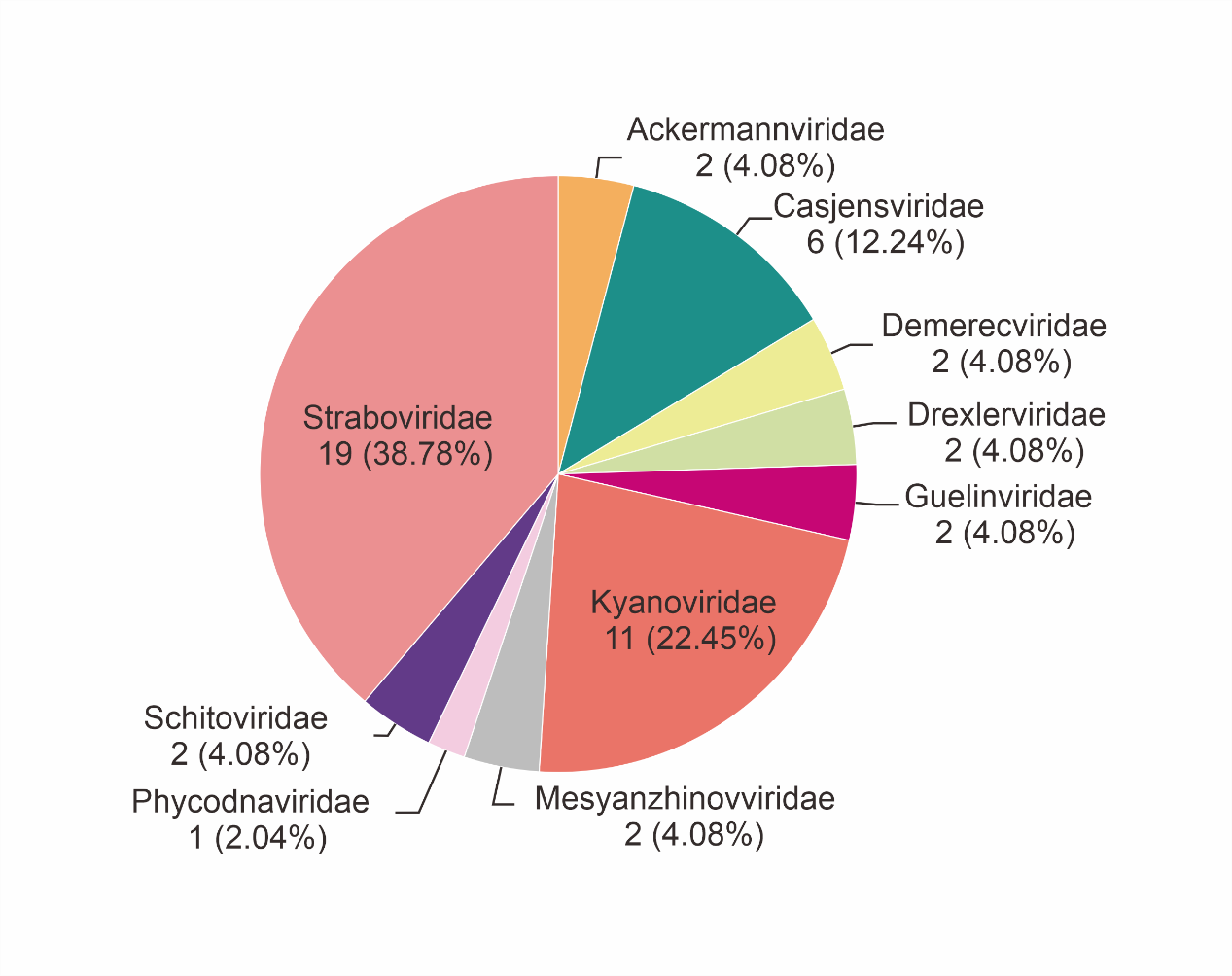

**Figure S15**. The family-level classifications of the phosphorus-cycling vContigs according to PhaGCN.
